# Activated ADAM10-targeting antibody 1H5 suppresses NOTCH-driven glioma progenitors

**DOI:** 10.64898/2026.10.28.762427

**Authors:** Umberto Tosi, Greta Ghita, Nayanendu Saha, Zaki Abou Mrad, Alexandra Giantini-Larsen, Kenny Yu, Johannes Jungverdorben, Dimitar B. Nikolov, Viviane Tabar, Yanhong Yang

## Abstract

**Background:** High-grade gliomas (HGGs) remain associated with high morbidity, mortality, and invariable recurrence after neurosurgical resection and chemoradiation. A major contributor to treatment failure is a population of stem-like glioma progenitor cells (GPCs) that depend on NOTCH-signaling for self-renewal and survival; clinical attempts to inhibit NOTCH with γ-secretase inhibitors failed due to systemic toxicity.

**Methods:** Here, we investigate the anti-HGG therapeutic potential of a fully-human monoclonal antibody, 1H5, that selectively targets activated-ADAM10, an α-secretase required for NOTCH receptor processing, with high tumor specificity. We show that ADAM10 is markedly overexpressed and highly activated in primary and recurrent HGGs, using patient tissue, RNA-sequencing data, and early-passage HGG cell lines.

**Results:** Across a panel of patient-derived HGG lines, sensitivity to 1H5 correlates with the cell-surface levels of activated-ADAM10, and shRNA-mediated ADAM10 knockdown recapitulates the growth-inhibitory effects of 1H5. ADAM10 inhibition suppresses NOTCH-pathway components and target genes, reduces expression of GPC markers, and impairs neurosphere formation while depleting Sox2- and Ki67-positive cells. We further find that 1H5 induces HGG cell death predominantly via necroptosis, characterized by survivin loss and increased phosphorylated MLKL and by transcriptional changes in necroptosis-related genes. In three-dimensional explants derived from primary HGG, 1H5 reduces growth and GPC marker expression while increasing TUNEL staining. Finally, convection-enhanced delivery of 1H5 in orthotopic xenograft models achieves high local intracranial concentrations, slows tumor growth, and prolongs survival, with regional pharmacodynamic effects consistent with NOTCH pathway suppression and necroptosis.

**Conclusions:** These data support ADAM10-targeting 1H5 as a candidate for further preclinical development in HGG.

## BACKGROUND

High-grade gliomas (HGGs) remain among the most lethal human cancers, with persistent high morbidity and near-universal mortality despite significant advances in the treatment of other solid and hematologic malignancies (1). Current standard therapy for glioblastoma multiforme (GBM), the most prevalent and aggressive subtype of HGG, consists of maximally safe surgical resection followed by radiotherapy and temozolomide chemotherapy. While this regimen modestly extends survival, median overall survival remains approximately 15 months, and essentially all tumors recur (2).

The ability of HGGs to survive therapy and recur has been linked to extensive cellular, spatial, and temporal heterogeneity. Single-cell and spatial profiling studies have revealed multiple co-existing cell states and microenvironmental niches, including perivascular, hypoxic, and invasive regions, each associated with distinct signaling dependencies and therapeutic sensitivities (3, 4). Over time, these heterogeneous compartments evolve under treatment pressure, giving rise to recurrent tumors that can differ substantially from their primaries in genetic composition, transcriptional state, and drug responses (5). A key contributor to this resilience is a population of stem-like glioma progenitor cells (GPCs), often marked by expression of SOX2, Nestin, and related stem cell factors, which exhibit enhanced self-renewal, treatment resistance, and capacity to repopulate the tumor after cytotoxic therapy. The presence and functional importance of such GPCs have been demonstrated in murine models, patient-derived cultures, and xenografts (6). Preclinical efforts at targeting GPCs focused on targeting NOTCH signaling, believed to be a key pathway in gliomagenesis (7).

NOTCH signaling is an evolutionarily conserved pathway that regulates cell fate, tissue development, and stem cell maintenance (8). NOTCH receptors comprise large extracellular epidermal growth factor (EGF)-like domains linked to transmembrane and intracellular domains (9). Ligand binding to the extracellular domain by members of the Jagged and Delta-like families triggers a two-step proteolytic cleavage, first by α-secretases of the ADAM (A Disintegrin and Metalloprotease) family and then by γ-secretase. These events release the NOTCH intracellular domain (NICD), which translocates to the nucleus and forms complexes with CSL (CBF1/RBPJκ, Suppressor of Hairless, Lag-1) transcription factors and co-regulators. The resulting transcriptional program promotes cell cycle progression, maintenance of a stem-like state, and, in appropriate contexts, tumorigenesis, with canonical targets including members of the HES and HEY helix-loop-helix families (10).

Given the central role of NOTCH signaling in GPC survival and function, multiple efforts have sought to target this pathway in GBM and other HGGs. Initial strategies focused on γ-secretase inhibitors such as RO4929097, which effectively downregulated NOTCH signaling in clinical trials based on downstream effector measurements but did not yield substantial clinical benefit due to dose-limiting systemic toxicities (11–15). Subsequent approaches turned to α-secretases, aiming to exploit their extracellular domains for antibody-mediated inhibition. Although preclinical studies demonstrated NOTCH pathway modulation with α-secretase targeting, translation remained limited, in part because of the broad physiological roles of these enzymes (16, 17).

ADAM10, a major α-secretase, has emerged as a candidate NOTCH regulator in GBM. Prior work has shown that ADAM10 is overexpressed in high-grade gliomas in a grade-dependent manner and that higher levels correlate with poorer prognosis (18). Importantly, ADAM10 exhibits lower expression across many normal tissues than γ-secretase components, suggesting that selective ADAM10 targeting could achieve a more favorable therapeutic window than global NOTCH blockade.

On glioma cells, ADAM10 adopts an activated, tumor-specific conformation and regulates cancer stem-like cells and tumor growth (19). This tumor-specific activated state was demonstrated to be ligand-independent and to promote tumorigenesis. Structural studies have further enabled development of monoclonal antibodies that recognize activated ADAM10 conformations through non-catalytic domains, providing a means to preferentially target tumor-associated ADAM10 activity. Consequently, targeting this activated state is expected to confer a high degree of tumor specificity, as autoinhibited ADAM10 in non-tumor tissues would be spared by activity-dependent therapeutic strategies.

In this study, we investigate ADAM10 as a therapeutic target in HGG and evaluate the efficacy of the fully human monoclonal antibody, 1H5, that binds the activated conformation of ADAM10 (20, 21). We characterize ADAM10 expression and activity in primary and recurrent GBM and other brain tumors, assess the relationship between activated ADAM10 levels and 1H5 sensitivity in patient-derived HGG cell lines, and examine the impact of ADAM10 inhibition on NOTCH signaling, GPC markers, self-renewal, and cell survival. We further delineate the mode of cell death induced by ADAM10 blockade, test 1H5 activity in a three-dimensional explant model derived from patient HGG tissue, and evaluate convection-enhanced delivery of 1H5 in orthotopic xenograft models. Together, these experiments aim to determine whether selective targeting of activated ADAM10 can disrupt NOTCH-dependent GPC maintenance and provide a foundation for developing ADAM10-directed antibodies as potential therapeutics for high-grade glioma.

## METHODS

### Cell Culture

Primary glioma samples from patients were harvested at the time of surgery then dissociated and cultured under N2 medium (DMEM/F12 catalog #11330032, Gibco, Carlsbad, CA) and supplied with N2 supplement (catalog #17502048, Gibco, Carlsbad, CA), B27 (NeuroCult™ SM1 Without Vitamin A, catalog #05731, Stemcell Technologies, Vancouver, Canada) and growth factors (Recombinant Human EGF Protein, catalog #236EG, and Recombinant Human FGF basic/FGF2/bFGF, catalog #233FB, both from R&D Systems, Minneapolis, MN). All the plates and dishes used for cell culture were pre-coated with Poly-L-ornithine hydrobromide (PO) (catalog #P3655, MilliporeSigma, Burlington, MA), Fibronectin (FN) (catalog # 356008, Fisher Healthcare, Waltham, MA) and murine Laminin I (catalog # 3400-010-02, R&D Systems, Minneapolis, MN).

### Purification of the 1H5 IgG form Expi293F cells

Generation of the 1H5 mAb from a large naïve human phage display library has been described previously (20). The 1H5 IgG was purified from Expi293F cells. Briefly, Expi293F cells were grown in Expi293 expression medium at 37°C, 8% CO2 with shaking at 125 rpmb. On the day of transfection, the cell density was maintained at 3-5 x 10^6^ cells/ml and viability at >95%. Heavy and light chain constructs cloned in pCAT2 vector backbone were mixed in Opti-MEM with linear polyethylenimine (PEI), a cationic polymer that aids transfection. For a 100 ml culture of Expi293F cells, 150 μg of each plasmid and 300 μg of PEI were used. The DNA:PEI solutions were incubated for 20 minutes at room temperature and added to the cells. After 20 hrs of incubation, enhancer 1 (100 mM sterile-filtered valproic acid (catalog #P4543-25G, MilliporeSigma, Burlington, MA) and enhancer 2 (100 mM sterile-filtered sodium propionate (catalog #P1880-500G, MilliporeSigma, Burlington, MA) were added and the cells were grown for additional 48 hrs. The 1H5 IgG was purified from the culture supernatant using proteinA Sepharose chromatography (affinity-matrix for IgG) followed by size-exclusion chromatography on a SD-200 column. The 1H5 IgG eluted with a molecular mass of 150 kDa on the SD-200 column and the heavy chain and light chains migrated at 50 and 25 kDa respectively on SDS-PAGE under reducing conditions (20).

### Cell-based ELISA assays to measure the binding of the anti-ADAM10 mAbs 1H5 and mAb1427 to glioblastoma cancer cells

Cellular ELISA was performed to measure the binding of the anti-ADAM10 human mAb 1H5, relative to the binding of the commercial anti-ADAM10 mAb1427 (monoclonal mouse IgG2B Clone # 163003, R&D), to ADAM10 expressed on the cell surface of glioma cell lines as well as to HEK293 cells and HEK293 cells transfected with full-length human ADAM10 (20, 22). Briefly, 5×104 cells/well were immobilized on 96-well ELISA plates with 1% paraformaldehyde for 2hrs at 37 °C. The plate was washed thrice with PBS and blocked for 2 hrs at room temperature with 4% non-fat dry milk. The anti-ADAM10 mAbs were then added in varying concentrations. Mouse mAb conjugated to HRP and recognizing human IgG was used as a secondary antibody (Abcam, Cambridge, United Kingdom) to detect 1H5 binding to ADAM10 expressed on the cells. For the commercial mAb1427 recognizing human ADAM10, we used goat anti-mouse IgG (H+L) secondary antibody conjugated to HRP (Thermo Fischer Scientific, Waltham, MA). Color was developed using the TMB substrate kit (Thermo Fischer Scientific, Waltham, MA). The data were recorded at 450 nm.

### Immunofluorescence (IF) staining and Immunohistochemistry (IHC) staining

Cells were plated on precoated chamber slides (154941PK, Fisher Scientific, Pittsburgh, PA) or 48-well plates and, following 24-hour attachment, treated with therapeutic-containing media (at concentrations highlighted above) or vehicle. At experimental timepoint, cells were fixed with 1% paraformaldehyde (PFA) in PBS for 10 minutes, washed with PBS, incubated for 1 hour at room temperature in blocking buffer – 5% normal goat serum, 0.1% Triton-X100 (catalog # 93443, Sigma, Burlington, MA) in PBS; washed with PBS, and incubated overnight at 4°C in primary antibody solution (a 1:10 dilution of blocking buffer in PBS). Primary antibodies utilized are reported in Supplemental Table 2. Following overnight incubation, cells were washed twice with PBS, incubated at room temperature and protected from light in secondary antibody solution, washed with PBS once, and incubated in a 2 µg / mL DAPI solution for 15 minutes. Following a final PBS wash, slides were mounted with Aqua-Pol y/Mount (catalog # 18606-20, Polysciences, Warrington, PA) before using a cover slide.

Patient- and mouse-derived fixed tissues were sectioned at 30 µm using a cryostat and mounted on slides. Following at least overnight incubation at -80°C, sections were allowed to thaw, washed with PBS three times, incubated for 1 hour at room temperature in blocking buffer (5% normal goat serum, 0.3% Triton-X100 in PBS), washed twice with PBS, and incubated overnight at 4°C in primary antibody solution (in a 1:10 dilution of blocking buffer in PBS) (Supplementary materials). Following overnight incubation, slides were washed thrice with PBS, incubated at room temperature and protected from light in secondary antibody solution (Supplementary materials), washed with PBS thrice, and incubated in a 2 µg / mL DAPI solution for 15 minutes. Following a final PBS wash, slides were mounted with Aqua-Poly/Mount (18606-20, Polysciences, Warrington, PA) before using a cover slide.

For paraffin-embedded sections, the staining process was preceded by deparaffinization and an antigen retrieval step, where sections were incubated for 30 minutes in a solution of 1 : 10 of VisUCyte Antigen Retrieval Reagent (Catalog # VCTS023, R & D Systems, Minneapolis, MN) in ddH_2_O in a water bath at 95°C. Sections were then blocked and incubated with primary antibodies overnight at 4°C, followed by HRP-conjugated secondary antibodies and DAB detection. Slides were counterstained with hematoxylin and mounted for imaging.

Images were taken with a Zeiss microscope-camera system and processed with ZEN Microscopy Softwarre (Zeiss, Oberkochen, Germany). Images were quantified using the FIJI Software (NIH, Bethesda, MD). Briefly, depending on the staining, a threshold was created to exclude background and, if applicable, weakly positive cells; images were made binary and processed with the watershed function. The automated particle analyzer plugin was utilized to obtain cell numbers. Alternatively, cells were counted manually. Thresholds were maintained constant across quantifications. Areas and signal intensities were measured using the appropriate software plugin.

### Cell viability assay with IC₅₀ determination

Primary tumor-derived cells between passage 5 and passage 25 were used in this study. Cells were dissociated by Accutase (catalog # NC9464543, Fisher Scientific, Pittsburgh, PA) into single cell suspensions, counted, and plated at 5000 cells / well in 96-well plates. Following 24-hour attachment, cells were treated with increasing concentration of therapeutic or vehicle. At experimental timepoint, cell viability was measured with AlamarBlue™ Cell Viability Reagent (catalog # DAL1100, Thermo Fisher Scientific, Pittsburgh, PA) and normalized against control-treated cells.

### ADAM10 knockdown via shRNA

Cells were thawed, plated, and grown to ∼90% confluency in a coated (PO + Fibronectin & Laminin) 6-well plate with GBM media. Cells were dissociated and 500,000 cells per condition (shNC, shADAM10-1, shADAM10-2) were collected and incubated with 5 μL of lentivirus (catalog# TRCN0000006675, MilliporeSigma, Burlington, MA) in Eppendorf tubes for 2 hours at 37 °C, with tubes agitated every 30 minutes to ensure transduction. Transduced cells were then plated at 250,000 cells per well and grown for 2 days prior to beginning puromycin selection along with a non-transduced control. MSK-001 was cultured with 1 μg/mL puromycin in GBM media (stock puromycin 5 mg/mL) for 1 week prior to collection for RNA-sequencing, Alamar blue, IF staining, and protein extraction for Western blotting. MSK-005 was cultured with 1 μg/mL of puromycin for a week prior to collection.

### Bulk RNA-seq Analysis

Cells from shNC and shADAM10 conditions were collected and submitted to Plasmidsaurus (Louisville, KY) for RNA sequencing. The vendor performed sample processing, library preparation, sequencing, and standard bioinformatic analysis according to their established workflow. Sequencing results, including processed gene expression data, were provided for subsequent downstream analyses

### Western Blotting

Primary tumor-derived cells were plated in 6-well plates and, following 24-hour attachment, treated with therapeutic-containing media (at subtherapeutic 133nM concentrations in PBS) or control. At experimental timepoints, cells were counted and lysed in loading buffer (50 mM Tris-HCl, pH 6.8; 2% SDS, 10% glycerin, 0.01% bromophenol blue, 100mM DTT). Following 30-minute incubation on ice, the lysates were boiled for 10 minutes. Proteins were separated by electrophoresis on 12% Bis-Tris gel in SDS running buffer for 1.5–2 hours and transferred to a nitrocellulose membrane. Non-specific protein binding was prevented by blocking the membrane with 5% Blotting-Grade Blocker (Bio-Rad Laboratories, Hercules, CA) in TBS-T (0.1% Tween-20 in 1x Tris-buffered saline). The membrane was incubated at 4°C overnight in the blocking buffer with primary antibodies (Supplementary Materials). After four washes with TBS-T, the blot was incubated with species-specific secondary antibodies for Mouse (1:5000) or Rabbit (1:5000) at room temperature for 2 hours and followed by four washes with TBS-T. ECL Prime Western Blotting Detection Kit was used for detection according to the manufacturer’s instruction.

### qRT-PCR

Primary tumor-derived cells were plated in 6-well plates and, following 24-hour attachment, treated with therapeutic-containing media (at subtherapeutic 133nM concentrations in PBS) or control. At experimental timepoints, cells were harvest and total RNA was extracted by TRIZOL (15596-018, Invitrogen, Thermo Fischer Scientific, Waltham, MA). Reverse transcription to cDNA was performed with the Quantitec Reverse transcription kit (205313, QIAGEN, Hilden, Germany) in accordance with the manufacturer’s instructions. Quantitative RT-PCR was performed using the SYBR Green PCR Master Mix (Applied Biosystems, Thermo Fischer Scientific, Waltham, MA) with the Biorad CFX 96 Real Time System. Primers are highlighted in Supplementary Materials

### Sphere Formation Assays

Cells were plated in coated 6-well plates at very low concentrations (between 25 and 200 cells per well) and, following overnight attachment, treated with therapeutic-containing media (at concentrations highlighted above) or control. Treatment occurred twice weekly for three consecutive weeks. At timepoint, media was removed and spheres fixed with a solution of 10% methanol and 10% glacial acetic acid in double distilled water (ddH_2_O) for 15 minutes at room temperature. A 1% solution of crystal purple in ddH_2_O was applied for 15 minutes, and then washed four times with ddH_2_O. The plates were allowed to dry overnight, scanned, and the number of spheres per condition manually counted. All experiments were repeated in triplicates.

### Cell Cycle Analysis

Primary tumor-derived cells were plated in 6-well plates and, following 24-hour attachment, cells were sequentially treated so that endpoint could be reached on the same day for all treatment conditions. At endpoint, cells were dissociated by Accutase (catalog # NC9464543, Fisher Scientific, Pittsburgh, PA), collected, and resuspended in a 70% solution of ethanol at 4°C. Cells were rested overnight at -20°C; the ethanol was removed and the cells were washed with PBS. Cells were then re-suspended in a PBS solution of 100 µg/mL RNase A (Ambion, Thermo Fischer Scientific, Waltham, MA) and 50 µg/mL of Propidium Iodide (Invitrogen, Thermo Fischer Scientific, Waltham, MA). Following 1-hour incubation, cells were washed with PBS and run through a flow cytometer.

### Explant Preparation and Treatment

Organotypic explants were processed from fresh tumor samples, on ice, immediately post-resection. The tumor tissue was dissected into smaller pieces (∼6 mm in length). Multiple small sphere-like explants are then punched out of the tumor pieces using a sterile 2 mm biopsy punch (Miltex, Integra Lifesciences, Princeton, NJ). Small spherical depressions were made in a sterile parafilm square and 10 µL of growth factor reduced Matrigel® (Corning Inc., Corning, NY) was added to each depression before transferring one 4 mm^3^ explant into each well, followed quickly by another 20 µL of Matrigel®. The Matrigel-embedded explants were allowed to fully polymerize in a 37°C incubator for 30 minutes before being transferred into a 6-well plate at 4-5 explants per well with 3 mL of explant culture media: DMEM/F-12 media (Gibco, Carlsbad, CA), N-2 media supplement (Gibco, Carlsbad, CA), and 1x Penicillin-Streptomycin (Gibco, Carlsbad, CA). Culture media was changed thrice weekly. After one week of rest, explants media was replaced with therapeutic-containing media or control. In some cases, 1H5 or control were directly injected into explants via a 32G Hamilton syringe (Hamilton, Reno, NV). Explant pictures were taken thrice weekly with an Olympus (Olympus Life Science, Thermo Fischer Scientific, Waltham, MA), processed with cellSens™ software (Olympus Life Science, Thermo Fischer Scientific, Waltham, MA), and analyzed using the FIJI Software (NIH, Bethesda, MD).

At endpoint, explants were removed from the media-therapeutic solution, fixed overnight in 4% PFA, followed by at least overnight incubation in 70% ethanol. They were them embedded in paraffin, sectioned at 5 µm thickness, and stained as described above.

### 1H5 Coupling with 488

1H5 was coupled with the fluorophore 488 using a large protein labeling kit (catalog # A10235, Thermo Fisher Scientific, Waltham, MA) according to manufacturer’s specifications.

### Animal Studies

All animal experiments were done in accordance with protocols approved by our Institutional Animal Care and Use Committee (IACUC) and following NIH guidelines for animal welfare.

#### Tumor implantation

Cells were dissociated with Accutase and the number of live cells was counted by Acridine Orange (AO) and Propidium Iodide (PI) staining (Nexcelom Bioscience, Lawrence, MA). Cells were resuspended in fresh medium at a density of 100’000 cells / µL. 2 µL were stereotactically injected in the right striatum. Male and female mice were evenly assigned to each group. Coordinates of the injection site were 1.7 mm lateral and 0.5 mm anterior to bregma, with 3.0 mm depth from skull. Mice were carefully monitored for up to the endpoint as designed for each study. NOD/SCID mice were used. For in vivo imaging, cells were lentivirally labeled with luciferase and tumor size was measured by Bioluminescence Imaging (BLI) using IVIS Spectrum In Vivo Imaging System to confirm tumor implantation. Survival of mice, or reach of humane experimental endpoints, as defined by the protocol (e.g. weight loss of 10% or more), was evaluated by Kaplan-Meier analysis and p-values were calculated using a log-rank test.

#### Convection Enhanced Delivery

therapeutics were delivered intracranially via convection-enhanced delivery (CED). Briefly, following confirmation of tumor implantation via BLI imaging, mice underwent delivery twice weekly for two consecutive weeks. Following anesthesia induction, the fur was shaved, and mice were securely placed on the stereotactic frame. Skin prep was performed with consecutive washes of betadine and 70% ethanol. The skin was opened and a 32G needle connected to a 50 µL Hamilton (Hamilton Company, Reno, NV) lowered at a depth of 3.0 mm from the skull at the site of prior cell implantation, through the same burr hole. 10 µL of therapeutic or vehicle were delivered over 5 minutes. The needle was slowly retracted one minute after the end of the infusion. The skin was sutured closed and the mice allowed to recover according to protocol.

### Statistical Analysis

Survival of mice was evaluated by Kaplan-Meier analysis and p-values were calculated using a log-rank test. Independent variables are compared using a two-tailed T-Test or a one-or two-way analysis of variance (ANOVA), as appropriate. All p-values are two-sided with statistical significance evaluated at the 0.05 alpha level. All analyses were performed ad in GraphPad Prism version 8 or subsequent (GraphPad Software, San Diego, CA).

## RESULTS

### ADAM10 Expression and Inhibition

To validate the targeting of ADAM10, its expression was first queried via immunofluorescence (IF) staining in primary tumor samples obtained at time of surgical resection. We found ADAM10 to be markedly overexpressed in primary and recurrent IDH-wildtype HGG, in contrast to its relatively low expression in IDH mutant gliomas (p = 0.006) and ependymomas (p = 0.03) (Figure 1.A and Supplemental Figure 1.A). The higher level of ADAM10 expression observed in IDH-wildtype HGG relative to IDH mutant gliomas (oligodendroglioma and astrocytoma) was also corroborated by bulk RNA sequencing data from TCGA (Supplemental Figure 1.B). ADAM10 expression was further validated in our early passage patient derived HGG cell lines, with co expression of SOX2 confirmed in representative lines (Figure 1.B). Conversely, ADAM10 exhibited lower systemic expression across most tissues than γ secretase, a previously targeted NOTCH protease (Supplemental Figure 1.C), suggesting a potentially reduced risk of systemic toxicity upon ADAM10 blockade than that observed following γ secretase inhibition. To assess the therapeutic potential of ADAM10 blockade in glioma, we developed a panel of low passage high grade glioma cell lines derived from surgical tissue. Detailed mutational analysis is presented in Supplemental Table 1. 1H5 preferentially binds the activated conformation of ADAM10 as previously described (Supplemental Figure 2.A, 2.B) (20, 21). The levels of activated ADAM10 on the cell surface were evaluated by ELISA across a panel of glioma cell lines as previously described (20). Among the seven lines tested, four demonstrated high levels of activated ADAM10, one line (MSK 003) exhibited moderate levels of activated ADAM10, and two lines (MSK 069 and MSK 006) showed minimal levels of activated ADAM10, with HK cells serving as a negative control (Figure 1.C). We then exposed these cell lines to 1H5. Notably, treatment with 1H5 inhibited HGG cell proliferation in a manner proportional to activated ADAM10 levels (Figure 1.D). HGG lines with high levels of activated ADAM10 displayed IC50 ranging from 110 to 800 nM, while the two lines with minimal activated ADAM10 levels only showed minimal reduction in viability within the same dose range (Figure 1.D). Human IgG was included as a background control and did not affect cell viability (Supplemental Figure 2.C). To further validate these findings, representative HGG lines were subjected to ADAM10 knockdown using shRNA. Suppression of ADAM10 expression resulted in a significant decrease in cell viability in two independent lines, an effect that was consistent with the reduction observed following 1H5 treatment (Figure 1.E and Supplemental Figure 3.A).

**Figure 1.**
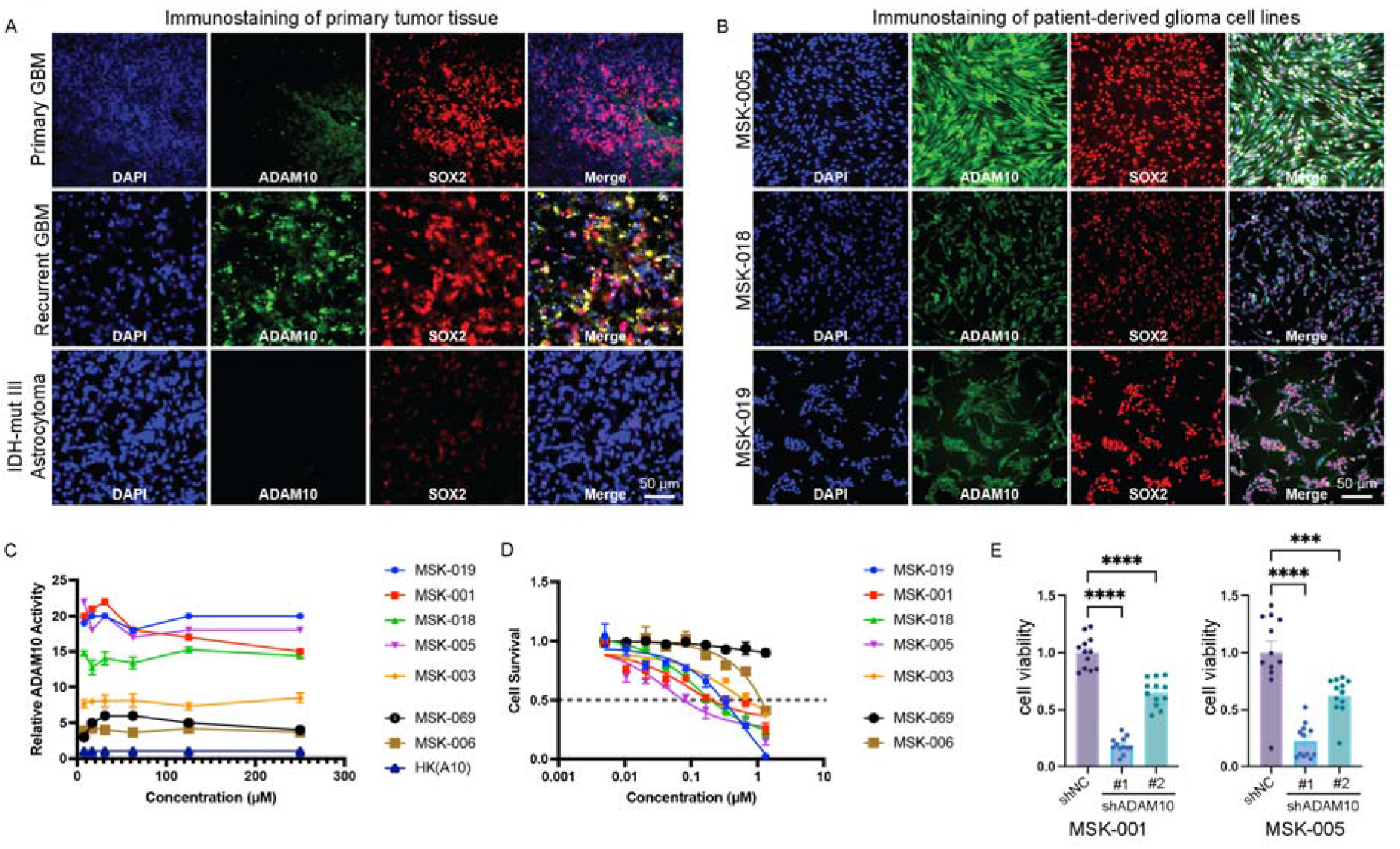
ADAM10 Expression and Inhibition. (A) immunofluorescence (IF) staining of tumor tissue shows ADAM10 and SOX2 (a marker for GPC) expression in primary and recurrent IDH-wildtype HGGs, with much lower expression in IDH-mutant samples. Scale bar, 50 μm. (B) IF of patient-derived cell lines showing high ADAM10 and SOX2 expression in patient-derived HGG lines. Scale bar, 50 μm. (C) ELISA-based quantification of relative level of activated ADAM10 across HGG cell lines. The assay identified cell lines with high levels of activated ADAM10, intermediate levels (MSK-003), and low levels of activated ADAM10. (D) IC50 curves (obtained via Alamar Blue assay) showing progressive response to increasing 1H5 dose following 6-day incubation; response to 1H5 is proportional to ADAM10 activity. (E) Cell viability is decreased following ADAM10 knockdown via shRNA across two HGG lines.

### 1H5 Efficacy via NOTCH Signaling Inhibition

To confirm whether the efficacy of 1H5 against HGGs is mediated through inhibition of the NOTCH signaling pathway, protein expression of the Notch pathway components was analyzed by western blotting. Treatment with 1H5 led to a marked reduction in ADAM10, the Notch1 receptor (N1R), the cleaved Notch receptor inner domain NICD, and the downstream effectors HES1, HES5, and HES6 (Figure 2.A). Consistent with these protein level changes, 1H5 treatment also significantly decreased the mRNA expression of the Notch target genes *HES1* and *HES6*(Figure 2.B and Supplementary Figure 3.B). To further validate these findings, RNA sequencing was performed in cells expressing shNC or shADAM10; ADAM10 knockdown resulted in significant global alterations in the transcriptional profile compared with controls (Figure 2.C). Pathway enrichment analysis performed using the Hallmark gene set revealed significant suppression of the NOTCH signaling pathway following shADAM10, while the TP53 signaling pathway was concomitantly activated (Figure 2.D). Consistent with these pathway-level changes, genes associated with the NOTCH pathway, including *ADAM10*, *NOTCH1*, *NOTCH2*, and *HES5*, as well as GPC markers such as *SOX2* and *NES*, were markedly downregulated (Figure 2.E). In contrast, TP53-associated genes, including *CDKN1A*, *CEBPA*, and *SERPINE1*, along with cell death–related genes such as *CABLES1*, were significantly upregulated following ADAM10 knockdown (Figure 2.E). Taken together, these results support that targeting ADAM10, through either antibody-mediated inhibition or shRNA knockdown, robustly suppresses the NOTCH signaling pathway, and glioma cell survival.

**Figure 2.**
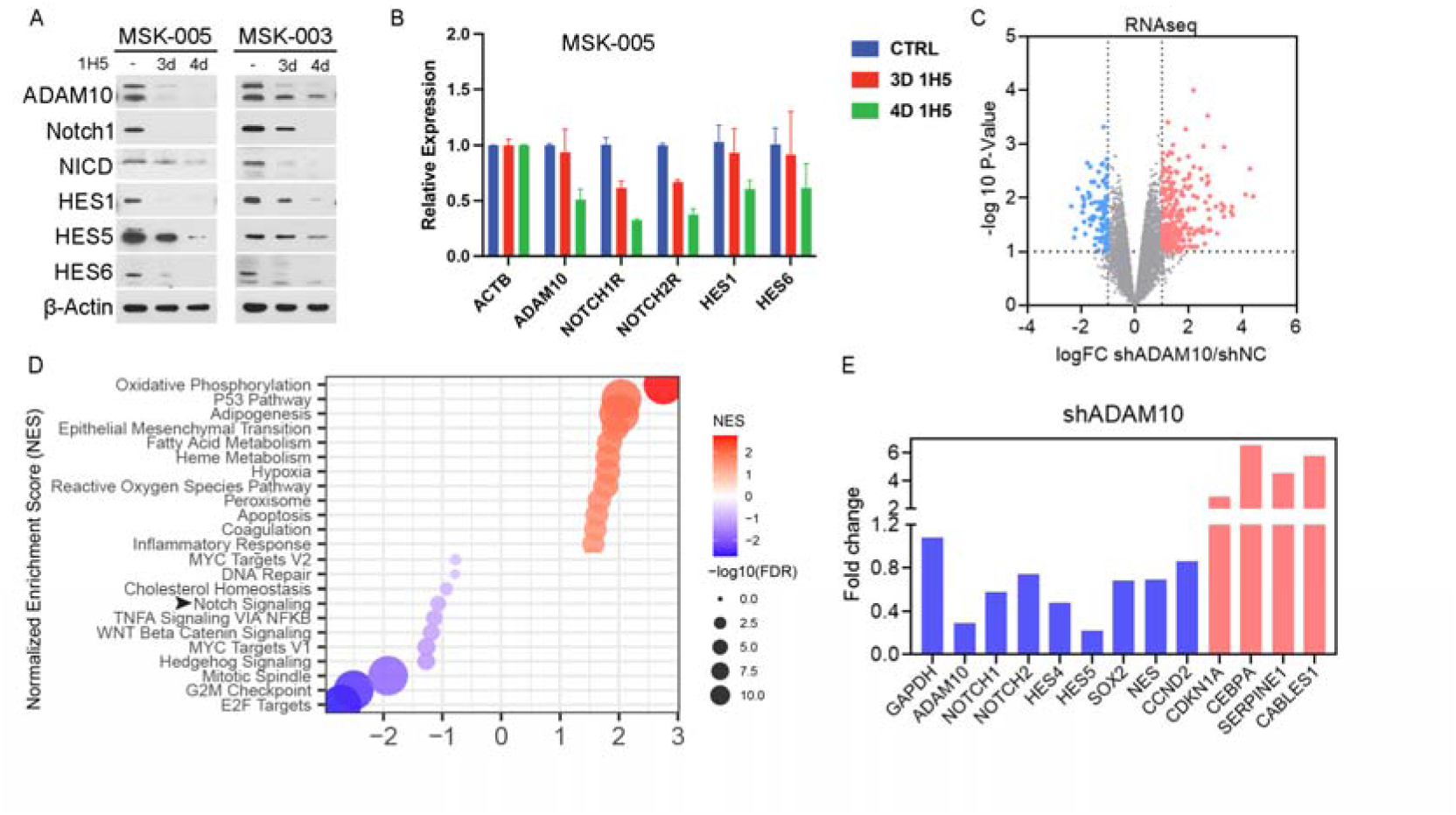
1H5 Efficacy via NOTCH Signaling Inhibition. (A, B) Western blotting (A) and qRT-PCR (B) performed following 1H5 treatment (133 nM) showing decrease in relevant NOTCH signaling pathway proteins and gene expression. (C, D) Bulk RNA-seq volcano plot (C) and pathway enrichment analysis (D) following ADAM10 knockdown via shRNA. (E) Gene expression following ADAM10 knockdown showing decrease in NOTCH-related genes (blue) and increase in TP53 associated genes (red).

### Targeting ADAM10 Reduces Self-Renewal Capacity of GPCs

Since the NOTCH signaling pathway plays a critical role in maintaining cellular self renewal in GPCs, tumorosphere formation assays were performed to assess whether 1H5 treatment would inhibit HGG cell self renewal. Treatment with 1H5 significantly reduced sphere formation in GBM cell lines compared to control (Figure 3.A, 3.B). Further, to determine whether 1H5 treatment depleted the GPC population, immunostaining was performed for the representative GPC marker SOX2 (Figure 3.C). A significant reduction in SOX2⁺ cells, proliferating Ki67⁺ cells, as well as double positive SOX2⁺/Ki67⁺ cells was observed following 1H5 treatment (Figure 3.D). The decrease in proliferating Ki67⁺ cells was further validated in two additional HGG cell lines (Figure 3.E).

**Figure 3.**
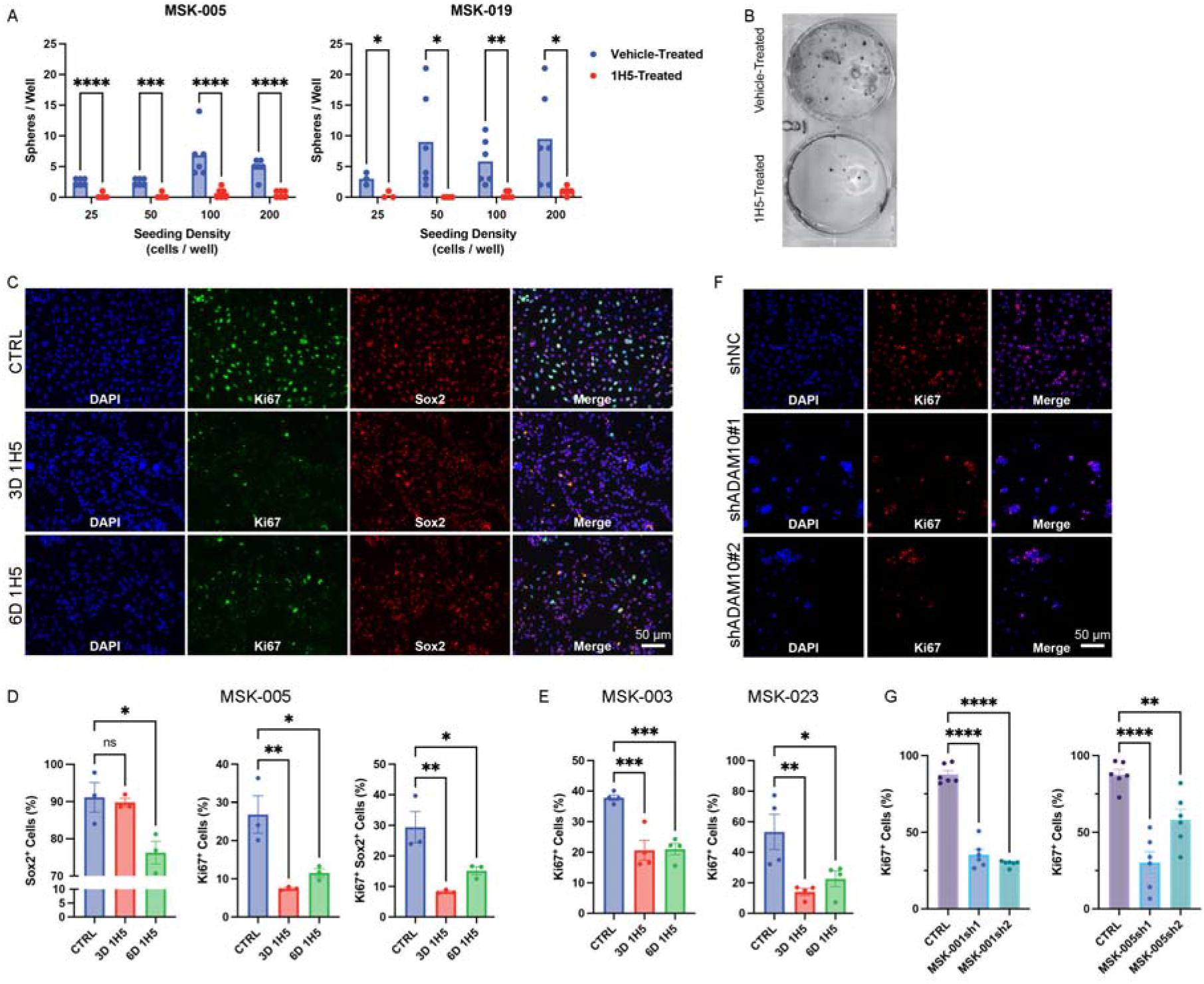
NOTCH Inhibition Results in Decreased Replicating Cells and GPCs. (A) 1H5 treatment (133 nM) reduces the ability to form neurospheres, a known marker of stemness, consistently and at increasing seeding densities. (B) Representative images of the neurosphere formation assay. Cells were seeded at a density of 100 cells per well and treated with either vehicle or 1H5 for 3 weeks, after which neurosphere formation was assessed. (C-E) IF (C) of cells treated with 1H5 (133nM for 3 or 6 days) shows a robust decrease in replicating (Ki67+) cells and SOX2+ GPCs, across multiple cell lines (D, E). Scale bar, 50 μm (F, G) IF of cells in which ADAM10 was knocked down (F) shows a significant decrease in replicating Ki67^+^ cells (G). Scale bar, 50 μm.

Consistent with the effects of 1H5 treatment, shRNA mediated knockdown of ADAM10 similarly resulted in a significant reduction of Ki67⁺ proliferating cells in HGG cell cultures (Figure 3.F, 3.G). Collectively, these results demonstrate that inhibition of ADAM10, either by 1H5 treatment or shRNA-mediated knockdown, significantly impairs HGG self renewal capacity and reduces the proliferative capacity of GPCs.

### 1H5 treatment triggers necroptosis-mediated cell death

We next investigated whether 1H5 exerted cytotoxic effects beyond suppressing self-renewal. HGG cells were treated with 1H5 and monitored over time. A significant reduction in Ki67^+^ cells was observed as early as day 2 following treatment and persisted through day 6 (Figure 4.A). Consistent with this finding, a time dependent decrease in total cell number was evident beginning on day 3 post-treatment. And numerous rounded and detached cells with a dying morphology were observed under brightfield microscopy on day 6 (Supplementary Figure 3.C). The dose-dependent decrease in cell viability and total cell number observed at day 6 was further validated in additional HGG cell lines (Supplementary Fig. 3D, 3.E).

**Figure 4.**
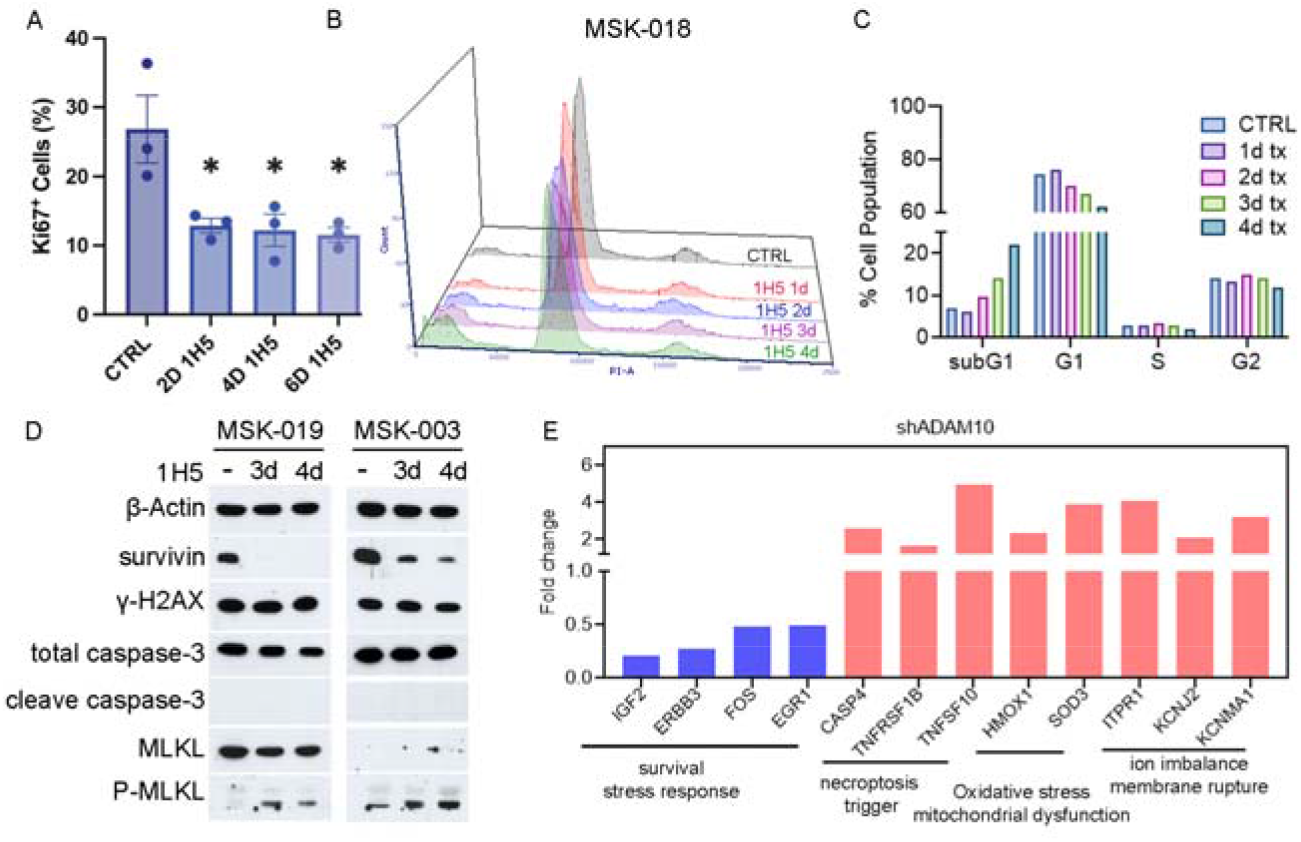
1H5 is cytotoxic causing cell necroptosis. (A) Treatment with 1H5 (300 nM) results in a decrease of replicating cells as early as two days following treatment initiation. (B, C) Flow cytometry for cell cycle analysis showing a progressive increase in a subG1 cell population indicating progressive cell death. (D) Western blotting following 1H5 treatment (133 nM) showing decrease in survivin no change in apoptosis markers γH2AX and cleaved caspase, with a robust increase in pMLKL, indicating necroptotic cell death. (E) RNA-seq following ADAM10 silencing showing decreased gene expression for survival response and increases in necroptosis, oxidative stress, and ion imbalance.

Cell death was further confirmed by flow cytometry, which revealed an increased sub-G1 population following 1H5 treatment (Figure 4.B, 4.C). This observation was independently validated in another HGG line, which similarly showed an elevated sub-G1 fraction at day 4 post-treatment (Supplementary Figure 3.F).

As accumulation of sub-G1 cells may result from apoptosis and/or necroptosis, we next examined key pathway markers by western blot. A marked reduction in survivin expression was detected, indicating impaired cell survival and progression toward cell death. However, no obvious increase in the DNA damage marker γH2AX or the apoptosis marker cleaved caspase-3 was observed, suggesting that apoptosis is not the primary mechanism of cell death. In contrast, we detected a robust increase in phosphorylated MLKL (p-MLKL), a hallmark of necroptosis (Figure 4.D). This finding was reproduced in additional HGG cell lines (Supplementary Figure 3.G).

In support of these results, RNA-seq analysis of shADAM10 cells revealed transcriptional changes consistent with necroptosis. Specifically, genes involved in cell survival and stress responses, including *IGF2*, *ERBB3*, *FOS*, and *EGR1*, were downregulated (Figure 2.D, 4E). In parallel, genes associated with necroptotic signaling (*TNFRSF1B*, *TNFSF10*), oxidative stress and mitochondrial dysfunction (*HMOX1*, *SOD3*), and ion imbalance and membrane rupture (*ITPR1*, *KCNJ2*, *KCNMA1*) were upregulated (Figure 4.E).

Collectively, these data indicate that blockade of ADAM10 by 1H5 induces HGG cell death predominantly through a necroptotic mechanism, similarly to ADAM10 downregulation via shRNA.

### 1H5 suppresses three-dimensional explant growth via reduction of replicating GPCs

To further evaluate the anti-glioma effects of 1H5, we employed an *ex vivo* glioma explant model. Organotypic three-dimensional explants are obtained by embedding 1 mm^3^ pieces of fresh surgical specimens in Matrigel. This approach maintains tumor composition and is known to replicate HGG heterogeneity and microenvironment *in vitro* (*7*). We generated explants from primary HGG tissue and treated longitudinally with 1H5 or vehicle (n = 8 per group). Explant growth area and radial growth distance were quantified as indicators of tissue viability and survival. As shown in Figure 5.A–5.C, 1H5 treatment resulted in a significant reduction in both growth area and growth distance compared with vehicle-treated explants. Both treated and control explants retained cytoarchitecture of the primary tumor (n = 8 per condition, Figure 5.D). Following treatment, explants were stained for relevant NOTCH-pathway proteins; 1H5-treated explants had a decrease in SOX2^+^ (p = 0.0037) and Ki67^+^ (p = 0.0064) cells (Figure 5 E, F) compared to vehicle treated controls. Similarly, a decrease in ADAM10^+^ (p = 0.0004) and Nestin^+^ (p = 0.0054) cells was observed (Figure 5.G, H). Consistent with *in vitro* evidence of 1H5 induced cell death and DNA fragmentation, as reflected by an increased sub G1 fraction, 1H5 treated explants exhibited elevated TUNEL staining, indicating increased DNA fragmentation in the explants (p < 0.0001, Figure 5.I, and Supplementary Figure 3.H). Taken together, these data suggest that 1H5 exhibits anti glioma effects in *ex vivo* models comparable to those observed in *in vitro* cell line studies.

**Figure 5.**
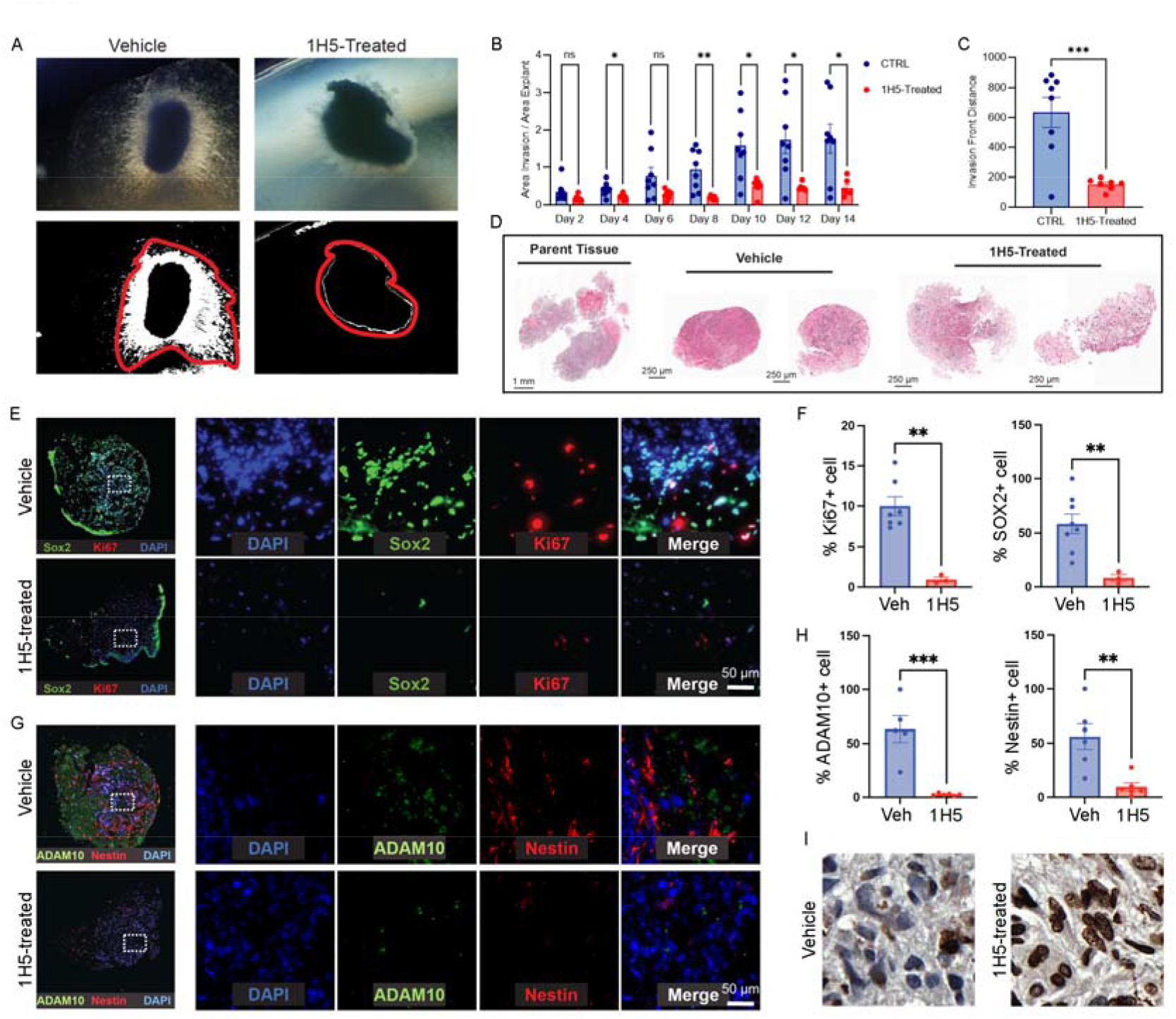
Three-dimensional HGG explants respond to 1H5 treatment. (A-C) Explants treated with 1H5 (266 nM) or vehicle were monitored for growth (A); 1H5 treatment resulted in decrease growth area (B) and maximal distance (C). Each data point represents one explant (n = 8 per group). (D) H&E staining of parent tumor and resultant explants demonstrate how three-dimensional explants retain numerous gross features of the original tumor. (E, F) IF (E) of vehicle-(top) and 1H5-treated (bottom) explants demonstrates a decrease in Ki67^+^ and SOX2^+^ cells. Scale bar, 50 μm. (G, H) IF (G) of vehicle-(top) and 1H5-treated (bottom) explants demonstrates a decrease in ADAM10^+^ and Nestin^+^ cells. Scale bar, 50 μm. (I) TUNEL staining shows increase in TUNEL signal in 1H5-treated explants over vehicle treated ones. Scale bar,

### CED of 1H5 prolongs survival of tumor-bearing mice

To characterize the *in vivo* behavior of 1H5, the antibody was first conjugated to a fluorophore for direct post-mortem visualization (“488-1H5”) and administered into the right striatum of animals via convection-enhanced delivery (CED), a technique where a catheter is placed in the target area in brain and used to deliver a drug solution under a pressure gradient to allow for coverage of large brain volumes with high infusate concentrations. To quantify intracranial 1H5 levels and estimate its half-life, animals receiving 488-1H5 via CED were sacrificed at progressive time points, ranging from immediately after infusion to 3 days post-CED. Brains were collected, and 1H5 levels were quantified based on the intensity of fluorescent signal. Separate cohorts received 488-1H5 via systemic administration (intraperitoneal injection) and were sacrificed 30 minutes later; these samples served as background controls.

CED delivery resulted in high intracranial concentrations of 488-1H5 (Supplemental Figure 4.A). Although antibody levels declined over the subsequent three days, brain concentrations at the final time point remained higher than those observed immediately following systemic administration (Supplemental Figure 4.B, 4.C). Fitting an exponential decay model to the clearance data yielded an estimated intracranial half-life of 1H5 of 17.6 hours (Supplemental Figure 4.C). Given the rapid clearance observed, where a significant portion of injected antibody was lost over three days, survival experiments were designed with CED repeated every fourth day, to maximize antibody efficacy while balancing procedural morbidity. The last CED was performed with 488-1H5 to obtain tumor coverage information (Figure 6.A). MSK-019 tumor implantation was confirmed via BLI; the animals were treated twice weekly via CED of 1H5 or vehicle over two weeks. Weight and animal behavior were monitored according to institutional protocols to confirm tolerability of the treatment. Following the last CED, tumor volume was again measured (Figure 6.B). The two groups had similar tumor volume before treatment (p = 0.96); following treatment, 1H5-treated mice had significantly smaller tumors that vehicle-treated counterparts (p = 0.034). Vehicle-treated tumors were significantly larger than pre-treatment (p = 0.0008) whereas 1H5-treated tumors were not (p = 0.07) (n = 11 per group, Figure 6.C). Similar results were obtained when mice transplanted with a slower-growing HGG line (MSK-001) were given time to recover following the third CED so that they could tolerate two more injections (Supplementary Figure 5.A). There were no significant weight differences between treatment and control groups (data not shown), and no obvious toxicity events were noted upon frequent mouse checks until significant tumor growth and animal demise. Importantly, 1H5-treated mice had a significantly longer survival than vehicle-treated controls (p = 0.013, Figure 6.D). Following the last CED, a small subset of animals was sacrificed and brains harvested for immunohistochemical analysis. Large right striatal tumors were observed in both 1H5- and vehicle-treated animals (Figure 6.E). In animals treated with 488-1H5, a region within the entire HNA^+^ tumor could be identified with strong 488^+^ signal, indicating higher 488-1H5 concentrations (Figure 6.F). Tumors were thus binned in regions with high and low 1H5 concentrations, which were then separated for analysis of downstream effectors (Figure 6.F). We observed that the tumor regions with high [1H5] had significantly lower levels of Ki67^+^ cells than regions with low [1H5] (p = 0.033) or vehicle-treated animals (p = 0.0006) (figure 6.G, Supplemental Figure 5.B); fewer SOX2^+^ cells (p < 0.0001 in both cases) (figure 6.H, Supplemental Figure 5.B); HES1^+^ cells (p < 0.0001 and p = 0.008, respectively) (figure 6.I, Supplemental Figure 5.C); and more pMLKL^+^ cells (p < 0.0001 for both groups) (figure 6.J, Supplemental Figure 5.D). This corroborates the *in vitro* finding of necroptosis as cell death mechanism following 1H5 treatment. Interestingly, ADAM10^+^ cells were lower in the high [1H5] areas than in vehicle treated animals (p < 0.0001) but not than in areas with low [1H5] (p = 0.13), which, however, had fewer ADAM10^+^ cells than vehicle (p < 0.0001) (Figure 6.K, Supplemental Figure 5.C). In summary, CED mediated delivery of 1H5 significantly suppressed tumor growth and extended animal survival, with mechanisms consistent with what was revealed by *in vitro* analyses.

**Figure 6.**
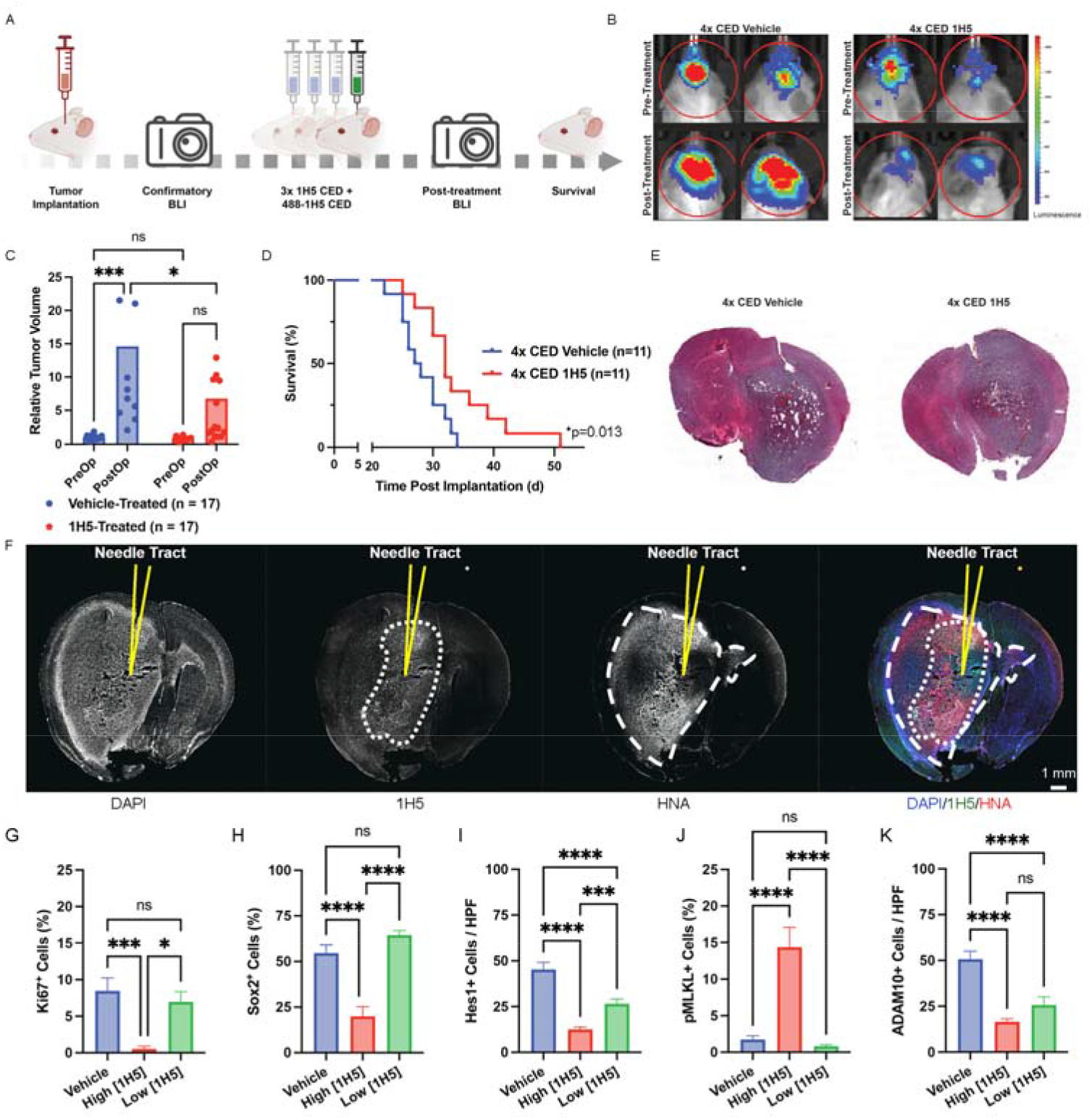
1H5 prolongs survival of HGG bearing mice via NOTCH inhibition. (A) Experimental schematics: following confirmation of tumor implantation, animals are injected with repeated CEDs of 1H5 or vehicle. Following the last CED, BLI imaging was performed. Created with BioRender.com. (B, C) Bioluminescence imaging (B) demonstrating a significantly slower tumor growth (MSK-019) in animals treated with CED of 1H5 compared to vehicle (C) (n = 17 per group). (D) Kaplan-Meyer curves demonstrating a significantly longer survival of animals treated with CED of 1H5 compared to vehicle (n = 11 per group). (E) H&E staining of animal brains treated with CED of 1H5 compared to vehicle demonstrating grossly similar tumor features. (F) Using CED of 488-1H5, an area within the tumor (HNA+ signal) with high 1H5 concentration was identified (“high [1H5]”) and compared to an area within the same tumor with low 1H5 concentration (“low [1H5]”). Scale bar, 1mm. (G-K) IF demonstrates significant differences across these two areas and vehicle-treated animals with respect to Ki67^+^ replicating cells (G), SOX2^+^ GPCs (H), HES1^+^ cells (I), pMLKL^+^ cells (J), and ADAM10^+^ cells (K).

## DISCUSSION

The present study identifies ADAM10 as a therapeutically relevant NOTCH regulator in high-grade glioma (HGG) and demonstrates that its selective targeting by the fully human monoclonal antibody 1H5 disrupts tumor growth through coordinated suppression of NOTCH signaling, depletion of glioma progenitor cells (GPCs), and induction of necroptotic cell death in vitro, ex vivo, and in vivo. Together, these findings integrate several converging themes in glioblastoma biology: the central importance of stem-like cell populations and intratumoral heterogeneity (*24–30*), the context-dependent role of NOTCH signaling in gliomagenesis (*6, 7, 31–34*), and the need for brain-penetrant biologics as tools to overcome the blood–brain barrier (*35–37*). By focusing on activated ADAM10 rather than global NOTCH inhibition, our data suggest a strategy that may combine effective suppression of a key stemness pathway with a potentially improved therapeutic window.

### ADAM10 as a context-specific NOTCH regulator in GBM

Our data show that ADAM10 is markedly overexpressed and functionally active in GBM relative to IDH-mutant gliomas and ependymomas, consistent with prior observations that proteolytic activation of NOTCH receptors is enriched in aggressive, stem-like tumor states (23, 24). Unlike broad γ-secretase inhibition, which has historically been limited by systemic toxicity and on-target gastrointestinal effects (11–15), ADAM10 appears to offer a more tumor-selective node of intervention. The relatively lower systemic expression of ADAM10 compared to γ-secretase supports the possibility of an improved therapeutic index.

Recent studies in HGG have further refined the understanding of NOTCH signaling as a dynamic regulator of cellular plasticity rather than a uniformly oncogenic pathway (25). NOTCH activity is now appreciated to fluctuate across transcriptional states (e.g., proneural-to-mesenchymal transitions) and to contribute to therapy resistance and recurrence (26, 27). Within this framework, our findings that 1H5 reduces NOTCH receptor activation (NICD), downstream effectors (HES family), and stemness markers (SOX2, Nestin) strongly support a model in which ADAM10 sustains a NOTCH-dependent progenitor state in GBM. The observed activation of TP53 signaling upon ADAM10 knockdown further suggests that NOTCH inhibition may relieve repression of tumor suppressor programs, aligning with emerging data that cross-talk between NOTCH and p53 pathways influences glioma cell fate decisions (28, 29).

### Targeting glioma progenitor cells and tumor self-renewal

Recent single-cell and lineage-tracing studies in GBM have emphasized that stemness is not a fixed property but a reversible state maintained by microenvironmental cues and signaling networks, including NOTCH (29–32). A key implication of these results is the selective vulnerability of GPCs to ADAM10 inhibition. GPCs are widely regarded as a reservoir for tumor propagation, therapeutic resistance, and recurrence in GBM (3–6). The marked reduction in neurosphere formation, SOX2+ populations, and proliferative marker Ki67 following 1H5 treatment indicates that ADAM10-NOTCH signaling is critical for maintaining this compartment.

### Induction of necroptosis as a therapeutic mechanism

Beyond its effects on proliferation and stemness, 1H5 induces robust cytotoxicity characterized by activation of necroptosis, as evidenced by increased phosphorylation of MLKL, TUNEL+ cells and transcriptional upregulation of necroptosis-associated genes. This is of particular importance given that GBM frequently exhibits resistance to apoptosis through alterations in pathways such as TP53, BCL-2 family signaling, and caspase activation (33).

Necroptosis, a regulated form of inflammatory cell death, has recently gained attention as a potential mechanism to overcome apoptosis resistance in cancer (34–36). In GBM, induction of necroptosis may have dual benefits: direct tumor cell killing and modulation of the tumor immune microenvironment through release of damage-associated molecular patterns (DAMPs). While our study did not directly assess immune consequences, this mechanism raises the possibility that ADAM10 inhibition could synergize with immunotherapeutic strategies, an area of increasing interest in HGG research (37–39).

### Implications for stratification of high-grade glioma patients

We had previously demonstrated how ADAM10 targeting via 1H5 was effective in preclinical models of colorectal cancer (20); success mainly came from targeting ADAM10’s non-catalytic domain, thus limiting side effects and toxicities arising from catalytic domain targeting (40). In glioma, the antibody also recognizes ADAM10 in its activated conformation, which is mostly expressed by tumor cells, thus limiting its toxicity. The local delivery via CED also serves to further reduce systemic side effects. In our previous studies, the impact of 1H5 was enhanced by combinatorial therapies, a feature that could be exploited in HGG therapies as well. Our data also suggest that tumor tissue assessment for the activated form of ADAM10 via ELISA can serve as a screen for tumors that are likely to respond to ADAM10 inhibition via 1H5. This would allow for a bedside to bench to bedside approach where, following index surgery for resection, the tumor is analyzed for expression of activated ADAM10 via ELISA, and if identified to be responsive to 1H5, therapy will be initiated accordingly.

### Translational relevance: overcoming the blood–brain barrier

A major challenge in developing antibody-based therapies for GBM is effective delivery across the blood–brain barrier (BBB). In this study, convection-enhanced delivery (CED), a technique shown to be clinically safe for brain tumor infusion in pediatric patients (41), enabled high local concentrations of 1H5, resulting in significant local tumor growth suppression and prolonged survival *in vivo*. This technique is very limited in rodents due to the difficulty of performing more than one intratumoral drug injection at a time, and to the lack of imaging guidance that ensures proper placement of the catheter in the tumor core. In humans, image-guided surgery allows multiple cannula passes to optimize maximal coverage of tumor volumes, as well as repeated delivery. While CED delivery effectively bypasses the blood brain barrier, the antibody product has a relatively short intracranial half-life, thus requiring repeated administration which is also challenging in rodents since every administration requires full anesthesia. These factors underscore the limitations of available delivery approaches and probably negatively impacted the efficacy outcomes of 1H5 intratumoral therapy. Nonetheless the data show promise and significant suppression of tumor progenitors and proliferation in the areas of high drug content.

Recent advances in bispecific antibody engineering offer promising alternatives for systemic delivery. BBB-penetrant bispecifics have demonstrated the ability to shuttle therapeutic antibodies into the brain parenchyma. Novel “shuttles” that bind the transferrin receptor are being developed to increase CNS-levels of large-molecule and viral therapeutics in neurodegenerative disease models (42–44); CD98hc targeting has shown promise in achieving persistently high levels of intracranial monoclonal antibodies in preclinical models (45–47); and p-selectin was recently identified as a targetable epitope that allows for transcytosis of nanoparticles in a medulloblastoma model (48, 49). Incorporating ADAM10-targeting moieties into such bispecific formats could enable non-invasive delivery while maintaining tumor specificity. Moreover, conditionally active or affinity-tuned bispecifics may further enhance selective accumulation within tumor tissue, minimizing off-target effects (50).

Applying similar strategies to ADAM10 inhibition could represent a logical next step, particularly given the strong dependency of a subset of GBMs on ADAM10 activity demonstrated here.

### Limitations and future directions

Several limitations of the study should be taken into consideration. First, while our in vitro and ex vivo models capture key aspects of high grade glioma biology, they do not fully recapitulate the complexity of the tumor microenvironment, including immune and vascular components. Second, the reliance on CED limits immediate clinical translatability, although it provides proof-of-principle for target engagement. However, CED in human patients offers greater precision of administration though repeat administration over time is limited. Bispecific antibodies could offer a potential solution by coupling 1H5 to a blood brain penetrant or facilitating antibody such as transferrin or others that allow for systemic delivery. Third, the heterogeneity in the levels of activated ADAM10across cell lines suggests that patient selection will be critical; biomarkers of ADAM10 activation or NOTCH dependency may help identify responsive tumors and the ELISA assay for activated ADAM10 used in this study can serve as a starting point for establishing tissue-based inclusion criteria for patients eligible for treatment.

Future studies could explore combination strategies, particularly with radiotherapy, immune checkpoint inhibitors, or agents targeting complementary pathways such as PI3K/AKT or MAPK signaling. Additionally, investigating the immunologic consequences of necroptosis induction in vivo could provide insight into potential synergy with immunotherapies.

## CONCLUSIONS

In this study, a novel monoclonal antibody, 1H5, inhibitory of the activated form of ADAM10 found at high concentrations in HGGs, showed significant preclinical potential in inhibiting glioma growth, causing cell death via necroptosis, through NOTCH-signaling inhibition that led to depletion of GPC populations. Further clinical studies are necessary for the expedited clinical translation of 1H5.

## LIST OF ABBREVIATIONS

ADAM: A Disintegrin and Metalloprotease
CED: Convection enhanced delivery
HGG: high grade glioma
GMB: glioblastoma multiforme
GPC: glioma progenitor cell
NICD: NOTCH intracellular domain
NOTCH: Neurogenic locus notch homolog protein

## DECLARATIONS

### Ethics approval and consent to participate

All experiments were conducted following institutional policies, protocols, and federal regulations. Human tissue was obtained and processed according to Memorial Sloan Kettering Protocol 03-12-019. All animal studies were performed following Ethics and IACUC approval of Protocol 03-12-019, in accordance with the protocol as detailed above and with institutional policies, federal regulations, and ethical standards.

### Consent to participate

not applicable.

### Consent for publication

Not applicable.

### Availability of data and materials

All data supporting the findings of this study are available within the paper and its Supplementary Information.

### Authors’ contributions

UT, GG, NS, AGL, ZAM: data collection

UT, GG, YY: figure generation

UT, YY, VT: manuscript preparation and revision

KY, JJ, DN, VT: experimental design

KY, JJ, DN, VT: support

All authors: approval of submitted manuscript

## Acknowledgements

N/A

## Conflict of Interest

the authors have no conflict of interest, financial or otherwise, to disclose.

**Supplemental Table 1.** List of cell lines used and their respective mutations, as determined by MSK IMPACT®.

**Supplemental Table 2.** Detailed information of the antibodies utilized.

## Data Availability Statement

The data generated in this study are available within the article and its supplementary data files, or upon request to the corresponding author.

## FIGURE LEGEND

**Supplemental Figure 1. Expression of ADAM10.** (A, B) Immunofluorescence staining (IF) shows ADAM10 and SOX2 (a marker for GPC) expression in Primary (n = 6) and recurrent (n = 4) HGG, with much lower expression in IDH-mutant samples (n = 5) or ependymomas (n = 2), as quantified (B). (C) ADAM10 mRNA expression in different glioma types, as determined via query of cBioPortal, normalized to normal brain, demonstrates preferentially higher expression of ADAM10 in higher grade samples. (D) Query of the gene expression atlas demonstrates significantly lower expression of ADAM10 protein systemically compared to γ-secretase.

**Supplemental Figure 2. 1 H5 Structure and ADAM10 inhibition.** (A) Superimpositions of the 1H5 Fab-bound ADAM10 D+C (pdb id: 8GH4) with a closed (pdb id: 6BE6) (left panel) and an open (pdb id; 8ESV) (right panel) conformations of the ADAM10 extracellular region. The ADAM10 proteinase domain is in red and the D+C domain region is in green. 1H5 is in cyan. The proteinase active site motif HExxH is shown as yellow spheres. (B) Schematics showing ADAM10 activation and mechanism of ADAM10 inhibition following 1H5 binding. (C) IC50 curves of cell lines treated with IgG control – no significant cell death is detected at 3 and 6 days.

**Supplemental Figure 3. ADAM10 Inhibition results in cell death via NOTCH signaling inhibition.** (A) Western blotting following shRNA knockdown of ADAM10 showing expected ADAM10 decrease in two HGG cell lines. (B) qRT-PCR performed following 1H5 treatment (133 nM) showing decrease in relevant NOTCH signaling pathway genes. (C) Brightfield microscopy showing progressive cell death following 1H5 treatment; by 6 days, cells detach and acquire a rounded, dying morphology. (D) Quantification of cell viability following treatment at increasing concentration of 1H5. (E) Immunofluorescence of cells treated with vehicle or 1H5 for 6 days showing significant decrease in live cells following 1H5 treatment. (F) Flow cytometry for cell cycle analysis showing a progressive increase in a subG1 cell population indicating progressive cell death. (G) Western blotting following 1H5 treatment (133 nM) showing decrease in survivin, no change in apoptosis markers γH2A.X and cleaved caspase 3, with a robust increase in pMLKL, indicating necroptotic cell death. (H) Explants treated with 1H5 (266 nM) have significantly higher proportion of TUNEL+ cells against vehicle-treated controls (p < 0.0001).

**Supplemental Figure 4. Fluorophore-labeled 1H5 allows for in vivo tracking following delivery.** (A) Following labeling of 488-1H5, it was delivered via CED in the right striatum of naïve mice, demonstrating progressive clearance over time. (B) When 488-1H5 was delivered systemically (IP), minimal signal was observed in the brain. (C) Quantification of signal intensity demonstrates an 8-fold increase in signal in the brain of mice injected with 488-1H5 via CED. A one-phase decay curve fitted to the data showed a clearance half-life of 17.6 hours.

**Supplemental Figure 5. CED of 1H5 Slows Down Tumor Growth via NOTCH Inhibition in PDX Models of HGGs.** (A) Quantification of BLI signal demonstrating a significantly slower tumor growth in animals implanted with MSK-001 treated with multiple CEDs of 1H5 (n = 12) compared to vehicle (n = 11). (B-D) Immunofluorescence imaging showing a decrease in Ki67^+^, SOX2^+^ (B), ADAM10^+^, HES1^+^ (C), and increase in pMLKL+ cells (D) following 1H5 treatment over vehicle.

## REFERENCES

1. Schaff LR, Mellinghoff IK. Glioblastoma and Other Primary Brain Malignancies in Adults: A Review. Jama. 2023;329(7):574–87.

2. Stupp R, Taillibert S, Kanner AA, Kesari S, Steinberg DM, Toms SA, et al. Maintenance Therapy With Tumor-Treating Fields Plus Temozolomide vs Temozolomide Alone for Glioblastoma: A Randomized Clinical Trial. Jama. 2015;314(23):2535–43.

3. Neftel C, Laffy J, Filbin MG, Hara T, Shore ME, Rahme GJ, et al. An Integrative Model of Cellular States, Plasticity, and Genetics for Glioblastoma. Cell. 2019;178(4):835–49.e21.

4. Ravi VM, Will P, Kueckelhaus J, Sun N, Joseph K, Salié H, et al. Spatially resolved multi-omics deciphers bidirectional tumor-host interdependence in glioblastoma. Cancer Cell. 2022;40(6):639–55.e13.

5. Kim J, Lee IH, Cho HJ, Park CK, Jung YS, Kim Y, et al. Spatiotemporal Evolution of the Primary Glioblastoma Genome. Cancer Cell. 2015;28(3):318–28.

6. Xie XP, Laks DR, Sun D, Ganbold M, Wang Z, Pedraza AM, et al. Quiescent human glioblastoma cancer stem cells drive tumor initiation, expansion, and recurrence following chemotherapy. Dev Cell. 2022;57(1):32–46.e8.

7. Hovinga KE, Shimizu F, Wang R, Panagiotakos G, Van Der Heijden M, Moayedpardazi H, et al. Inhibition of notch signaling in glioblastoma targets cancer stem cells via an endothelial cell intermediate. Stem Cells. 2010;28(6):1019–29.

8. Kopan R, Ilagan MX. The canonical Notch signaling pathway: unfolding the activation mechanism. Cell. 2009;137(2):216–33.

9. Li X, Yan X, Wang Y, Kaur B, Han H, Yu J. The Notch signaling pathway: a potential target for cancer immunotherapy. J Hematol Oncol. 2023;16(1):45.

10. Borggrefe T, Oswald F. The Notch signaling pathway: transcriptional regulation at Notch target genes. Cell Mol Life Sci. 2009;66(10):1631–46.

11. Dantas-Barbosa C, Bergthold G, Daudigeos-Dubus E, Blockus H, Boylan JF, Ferreira C, et al. Inhibition of the NOTCH pathway using γ-secretase inhibitor RO4929097 has limited antitumor activity in established glial tumors. Anticancer Drugs. 2015;26(3):272–83.

12. Luistro L, He W, Smith M, Packman K, Vilenchik M, Carvajal D, et al. Preclinical profile of a potent gamma-secretase inhibitor targeting notch signaling with in vivo efficacy and pharmacodynamic properties. Cancer Res. 2009;69(19):7672–80.

13. Olsauskas-Kuprys R, Zlobin A, Osipo C. Gamma secretase inhibitors of Notch signaling. Onco Targets Ther. 2013;6:943–55.

14. Xu R, Shimizu F, Hovinga K, Beal K, Karimi S, Droms L, et al. Molecular and Clinical Effects of Notch Inhibition in Glioma Patients: A Phase 0/I Trial. Clin Cancer Res. 2016;22(19):4786–96.

15. Peereboom DM, Ye X, Mikkelsen T, Lesser GJ, Lieberman FS, Robins HI, et al. A Phase II and Pharmacodynamic Trial of RO4929097 for Patients With Recurrent/Progressive Glioblastoma. Neurosurgery. 2021;88(2):246–51.

16. Fan X, Khaki L, Zhu TS, Soules ME, Talsma CE, Gul N, et al. NOTCH pathway blockade depletes CD133-positive glioblastoma cells and inhibits growth of tumor neurospheres and xenografts. Stem Cells. 2010;28(1):5–16.

17. Takebe N, Nguyen D, Yang SX. Targeting notch signaling pathway in cancer: clinical development advances and challenges. Pharmacol Ther. 2014;141(2):140–9.

18. Qu M, Qiu BO, Xiong W, Chen D, Wu A. Expression of a-disintegrin and metalloproteinase 10 correlates with grade of malignancy in human glioma. Oncol Lett. 2015;9(5):2157–62.

19. Atapattu L, Saha N, Chheang C, Eissman MF, Xu K, Vail ME, et al. An activated form of ADAM10 is tumor selective and regulates cancer stem-like cells and tumor growth. J Exp Med. 2016;213(9):1741–57.

20. Saha N, Baek DS, Mendoza RP, Robev D, Xu Y, Goldgur Y, et al. Fully human monoclonal antibody targeting activated ADAM10 on colorectal cancer cells. Biomed Pharmacother. 2023;161:114494.

21. Mashati P, Sun D, Reino EG, Militello G, Gao N, Blandino JA, et al. Inhibition of colorectal cancer progression through conformation-specific targeting of ADAM10 metalloprotease. Front Oncol. 2025;15:1704436.

22. Saha N, Xu K, Zhu Z, Robev D, Kalidindi T, Xu Y, et al. Inhibitory monoclonal antibody targeting ADAM17 expressed on cancer cells. Transl Oncol. 2022;15(1):101265.

23. Martínez-Gascueña P, Nueda ML, Baladrón V. Modulation of Oncogenic NOTCH Signaling in Highly Aggressive Malignancies by Targeting the γ-Secretase Complex: A Systematic Review. Cells. 2026;15(5).

24. Shi Q, Xue C, Zeng Y, Yuan X, Chu Q, Jiang S, et al. Notch signaling pathway in cancer: from mechanistic insights to targeted therapies. Signal Transduct Target Ther. 2024;9(1):128.

25. D’Amico M, De Amicis F. Aberrant Notch signaling in gliomas: a potential landscape of actionable converging targets for combination approach in therapies resistance. Cancer Drug Resist. 2022;5(4):939–53.

26. De Silva MI, Stringer BW, Bardy C. Neuronal and tumourigenic boundaries of glioblastoma plasticity. Trends Cancer. 2023;9(3):223–36.

27. Parmigiani E, Taylor V, Giachino C. Oncogenic and Tumor-Suppressive Functions of NOTCH Signaling in Glioma. Cells. 2020;9(10).

28. Chowdhury S, Sarkar RR. Exploring Notch Pathway to Elucidate Phenotypic Plasticity and Intra-tumor Heterogeneity in Gliomas. Sci Rep. 2019;9(1):9488.

29. Wang X, Zhou R, Xiong Y, Zhou L, Yan X, Wang M, et al. Sequential fate-switches in stem-like cells drive the tumorigenic trajectory from human neural stem cells to malignant glioma. Cell Res. 2021;31(6):684–702.

30. Eisenbarth D, Wang YA. Glioblastoma heterogeneity at single cell resolution. Oncogene. 2023;42(27):2155–65.

31. Patel AP, Tirosh I, Trombetta JJ, Shalek AK, Gillespie SM, Wakimoto H, et al. Single-cell RNA-seq highlights intratumoral heterogeneity in primary glioblastoma. Science. 2014;344(6190):1396–401.

32. Prager BC, Bhargava S, Mahadev V, Hubert CG, Rich JN. Glioblastoma Stem Cells: Driving Resilience through Chaos. Trends Cancer. 2020;6(3):223–35.

33. Guo J, Zong L, Huang Y, Liu X, Hu Y, Liu Y. Unlocking glioma vulnerabilities: targeting regulated cell death pathways for innovative therapies. Cell Death Discov. 2026;12(1):95.

34. Li Y, Qiu Y, Gao W, Geng L, Wu Z, Zhang T, et al. Necroptotic signaling orchestrates glioblastoma malignancy and potentiates temozolomide response. Cell Death Dis. 2025;16(1):921.

35. Pagano C, Navarra G, Coppola L, Avilia G, Pastorino O, Della Monica R, et al. N6-isopentenyladenosine induces cell death through necroptosis in human glioblastoma cells. Cell Death Discov. 2022;8(1):173.

36. Wu Y, Dong G, Sheng C. Targeting necroptosis in anticancer therapy: mechanisms and modulators. Acta Pharm Sin B. 2020;10(9):1601–18.

37. Khan M, Huang X, Ye X, Zhang D, Wang B, Xu A, et al. Necroptosis-based glioblastoma prognostic subtypes: implications for TME remodeling and therapy response. Ann Med. 2024;56(1):2405079.

38. Sprooten J, De Wijngaert P, Vanmeerbeerk I, Martin S, Vangheluwe P, Schlenner S, et al. Necroptosis in Immuno-Oncology and Cancer Immunotherapy. Cells. 2020;9(8).

39. Zhang Z, Zhang F, Xie W, Niu Y, Wang H, Li G, et al. Induced Necroptosis and Its Role in Cancer Immunotherapy. Int J Mol Sci. 2024;25(19).

40. Janes PW, Saha N, Barton WA, Kolev MV, Wimmer-Kleikamp SH, Nievergall E, et al. Adam meets Eph: an ADAM substrate recognition module acts as a molecular switch for ephrin cleavage in trans. Cell. 2005;123(2):291–304.

41. Souweidane MM, Kramer K, Pandit-Taskar N, Zhou Z, Haque S, Zanzonico P, et al. Convection-enhanced delivery for diffuse intrinsic pontine glioma: a single-centre, dose-escalation, phase 1 trial. Lancet Oncol. 2018;19(8):1040–50.

42. Barker SJ, Thayer MB, Kim C, Tatarakis D, Simon MJ, Dial R, et al. Targeting the transferrin receptor to transport antisense oligonucleotides across the mammalian blood-brain barrier. Sci Transl Med. 2024;16(760):eadi2245.

43. Huang Q, Chan KY, Wu J, Botticello-Romero NR, Zheng Q, Lou S, et al. An AAV capsid reprogrammed to bind human transferrin receptor mediates brain-wide gene delivery. Science. 2024;384(6701):1220–7.

44. Sonoda H, Morimoto H, Yoden E, Koshimura Y, Kinoshita M, Golovina G, et al. A Blood-Brain-Barrier-Penetrating Anti-human Transferrin Receptor Antibody Fusion Protein for Neuronopathic Mucopolysaccharidosis II. Mol Ther. 2018;26(5):1366–74.

45. Chew KS, Wells RC, Moshkforoush A, Chan D, Lechtenberg KJ, Tran HL, et al. CD98hc is a target for brain delivery of biotherapeutics. Nat Commun. 2023;14(1):5053.

46. Pornnoppadol G, Bond LG, Lucas MJ, Zupancic JM, Kuo YH, Zhang B, et al. Bispecific antibody shuttles targeting CD98hc mediate efficient and long-lived brain delivery of IgGs. Cell Chem Biol. 2024;31(2):361–72.e8.

47. Veire JL, Lucas MJ, Bond LG, Tripu DR, Tessier PM, Greineder CF. Comparative Radiotracing Quantifies Brain Cellular Uptake and Catabolism of Bispecific Antibodies Targeting Transferrin Receptor and CD98hc. ACS Chem Neurosci. 2025;16(7):1264–74.

48. Shamay Y, Elkabets M, Li H, Shah J, Brook S, Wang F, et al. P-selectin is a nanotherapeutic delivery target in the tumor microenvironment. Sci Transl Med. 2016;8(345):345ra87.

49. Tylawsky DE, Kiguchi H, Vaynshteyn J, Gerwin J, Shah J, Islam T, et al. P-selectin-targeted nanocarriers induce active crossing of the blood-brain barrier via caveolin-1-dependent transcytosis. Nat Mater. 2023;22(3):391–9.

50. Wells RC, Akkapeddi P, Chan D, Joy D, Moshkforoush A, Chau R, et al. Dual targeting of transferrin receptor and CD98hc enhances brain exposure of large molecules. Cell Rep. 2025;44(8):116038.

