## Supplementary_Figures&Tables for "Activated ADAM10-targeting antibody 1H5 suppresses NOTCH-driven glioma progenitors"

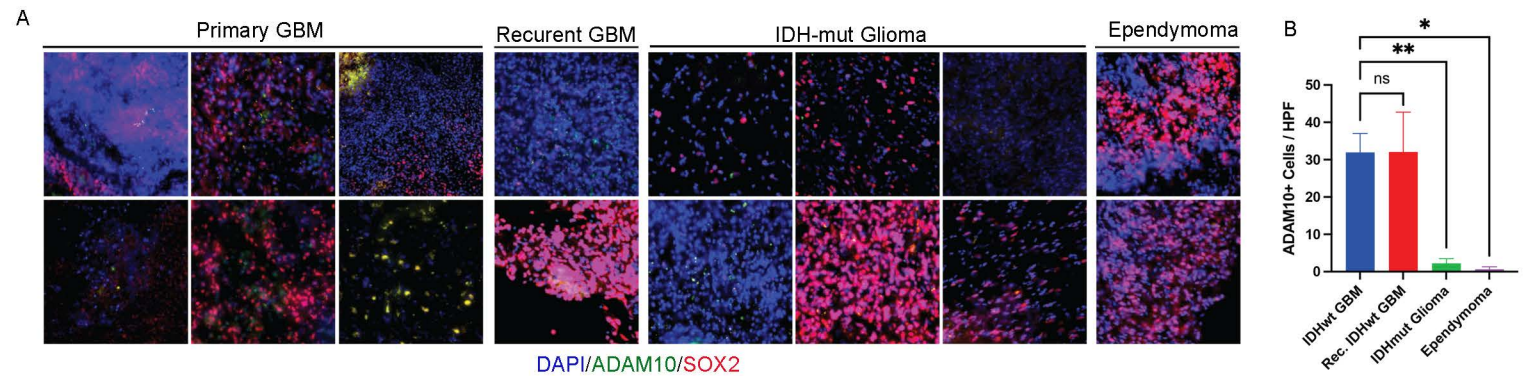

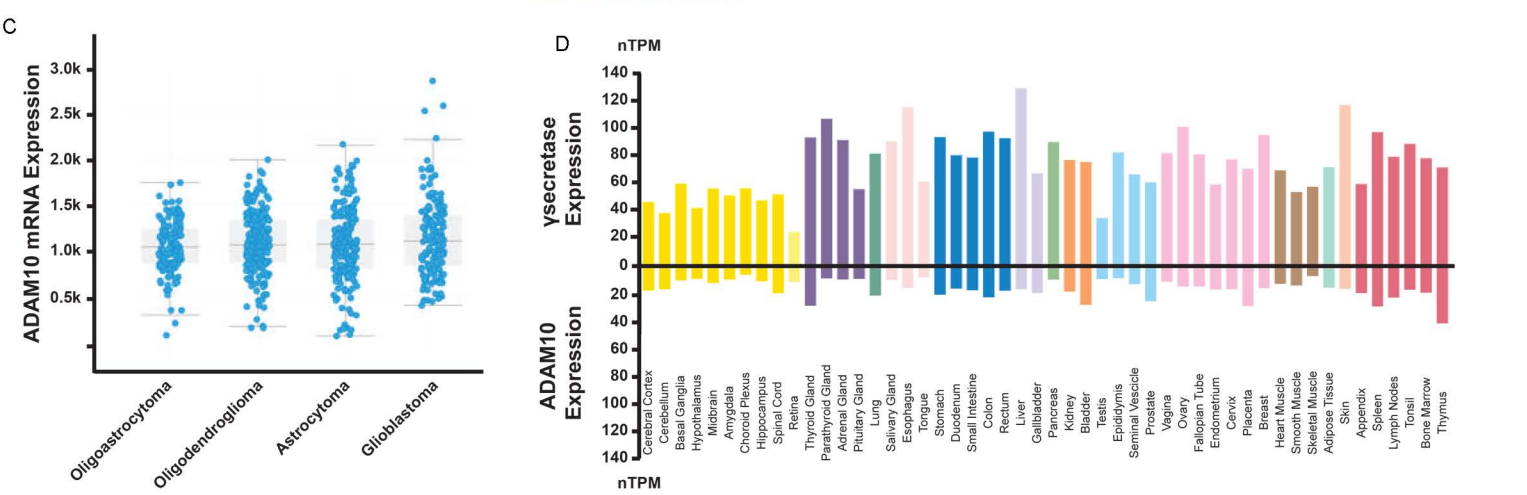

**Supplemental Figure 1. Expression of ADAM10.** (A, B) Immunofluorescence staining (IF) shows ADAM10 and SOX2 (a marker for GPC) expression in Primary (n = 6) and recurrent (n = 4) HGG, with much lower expression in IDH-mutant samples (n = 5) or ependymomas (n = 2), as quantified (B). (C) ADAM10 mRNA expression in different glioma types, as determined via query of cBioPortal, normalized to normal brain, demonstrates preferentially higher expression of ADAM10 in higher grade samples. (D) Query of the gene expression atlas demonstrates significantly lower expression of ADAM10 protein systemically compared to γ-secretase.

A

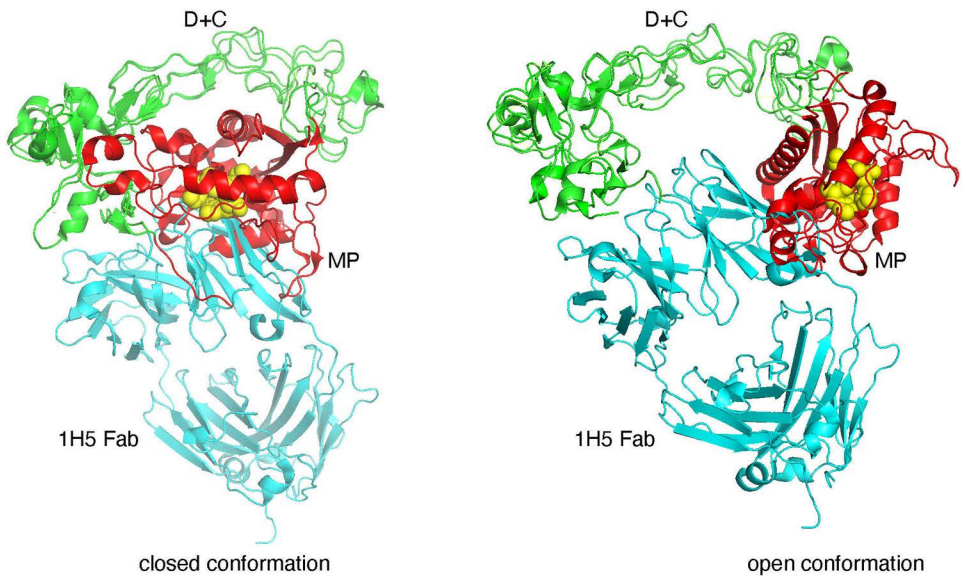

B

Schematic representation of a proposed mechanism for ADAM10 activation and interactions with substrates and the 1H5 antibody.

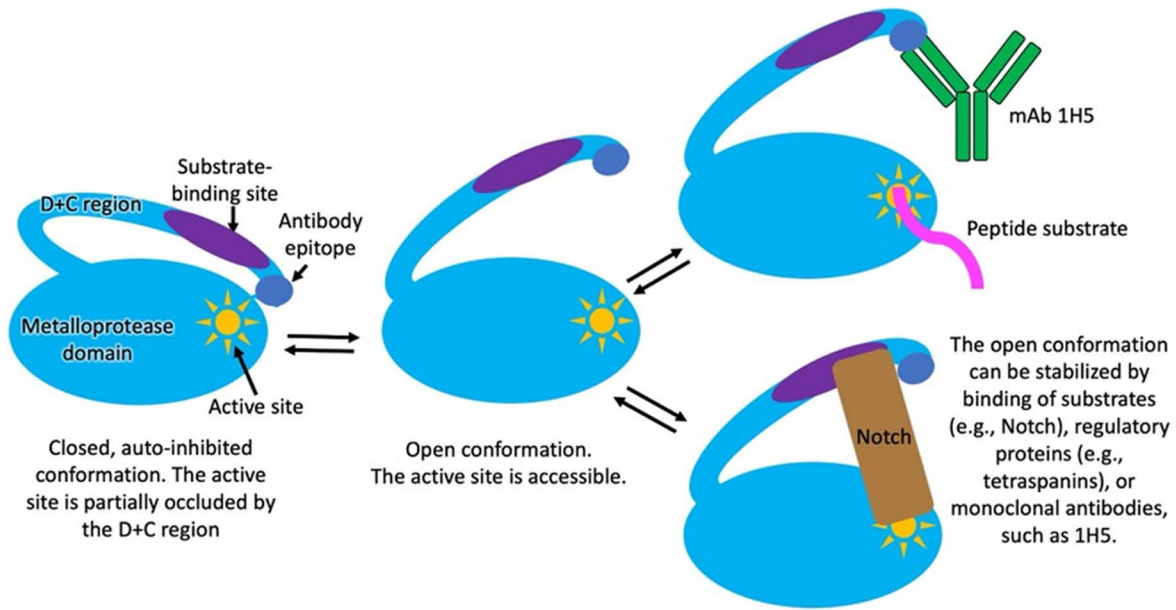

C

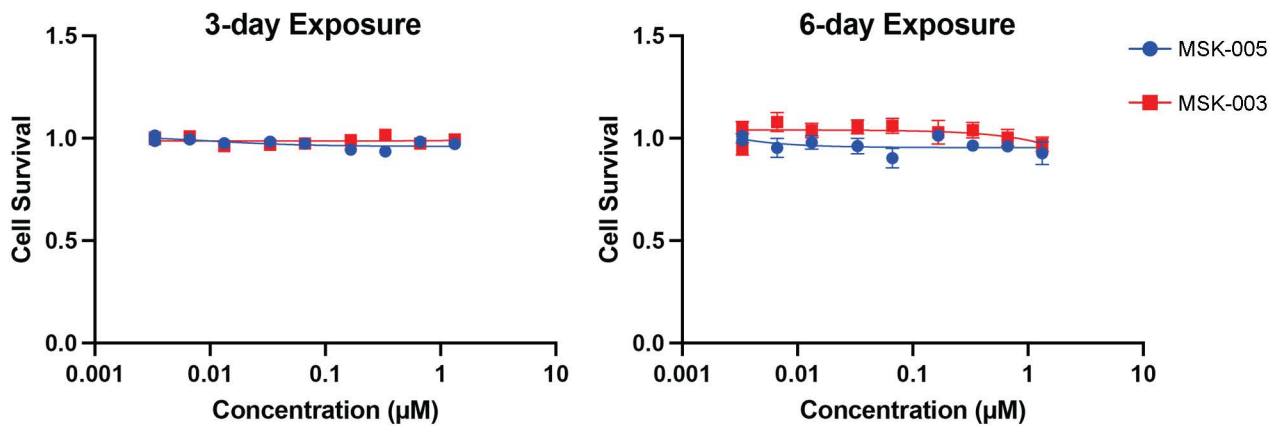

**Supplemental Figure 2. 1H5 Structure and ADAM10 inhibition.** (A) Superimpositions of the 1H5 Fab-bound ADAM10 D+C (pdb id: 8GH4) with a closed (pdb id: 6BE6) (left panel) and an open (pdb id; 8ESV) (right panel) conformations of the ADAM10 extracellular region. The ADAM10 proteinase domain is in red and the D+C domain region is in green. 1H5 is in cyan. The proteinase active site motif HExxH is shown as yellow spheres. (B) Schematics showing ADAM10 activation and mechanism of ADAM10 inhibition following 1H5 binding. (C) IC<sub>50</sub> curves of cell lines treated with IgG control – no significant cell death is detected at 3 and 6 days.

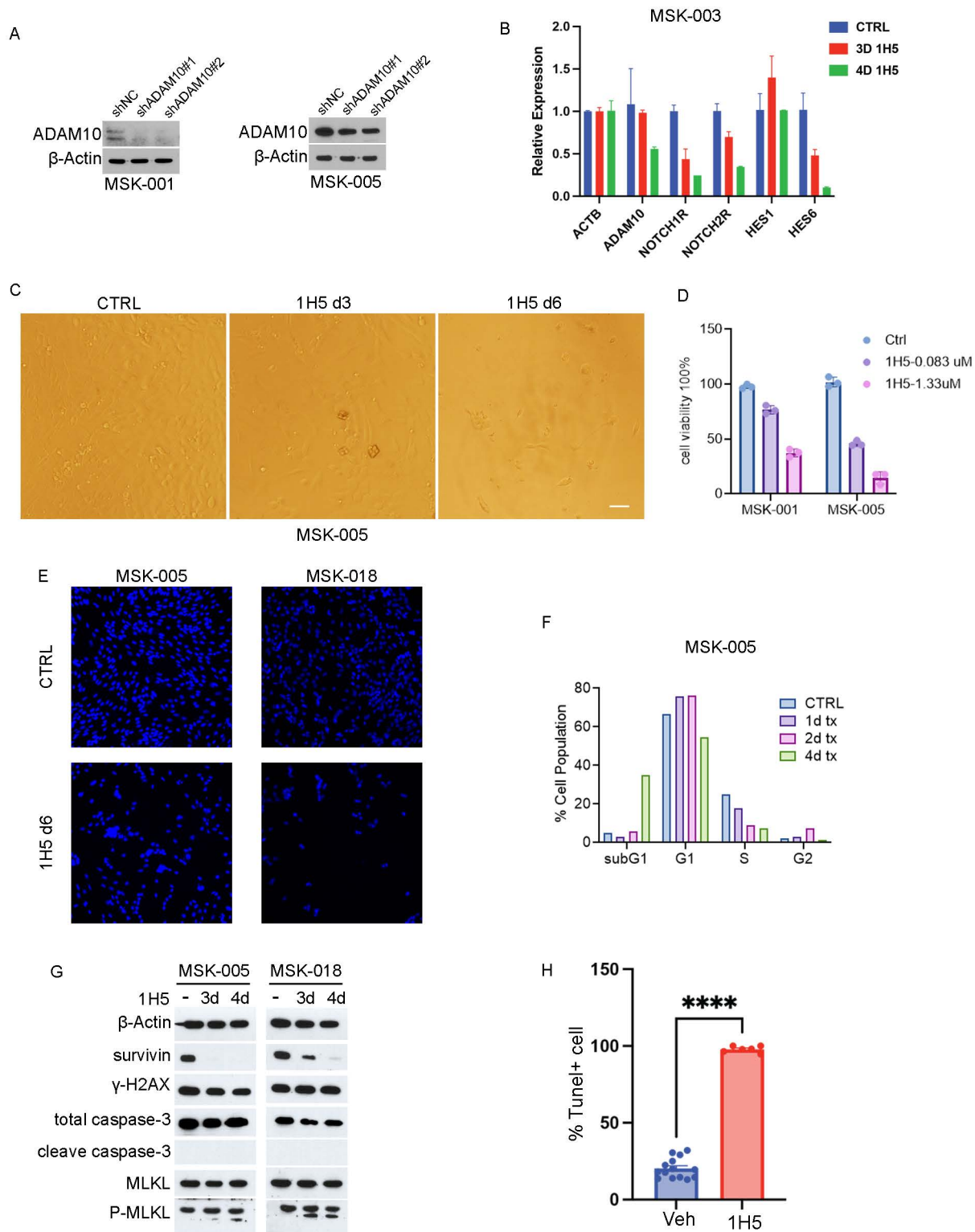

**Supplemental Figure 3. ADAM10 Inhibition results in cell death via NOTCH signaling inhibition.** (A) Western blotting following shRNA knockdown of ADAM10 showing expected ADAM10 decrease in two HGG cell lines. (B) qRT-PCR performed following 1H5 treatment (133 nM) showing decrease in relevant NOTCH signaling pathway genes. (C) Brightfield microscopy showing progressive cell death following 1H5 treatment; by 6 days, cells detach and acquire a rounded, dying morphology. (D) Quantification of cell viability following treatment at increasing concentration of 1H5. (E) Immunofluorescence of cells treated with vehicle or 1H5 for 6 days showing significant decrease in live cells following 1H5 treatment. (F) Flow cytometry for cell cycle analysis showing a progressive increase in a subG1 cell population indicating progressive cell death. (G) Western blotting following 1H5 treatment (133 nM) showing decrease in survivin, no change in apoptosis markers γH2A.X and cleaved caspase 3, with a robust increase in pMLKL, indicating necroptotic cell death. (H) Explants treated with 1H5 (266 nM) have significantly higher proportion of TUNEL+ cells against vehicle-treated controls ($p < 0.0001$).

**A**

**T = 0 minutes**

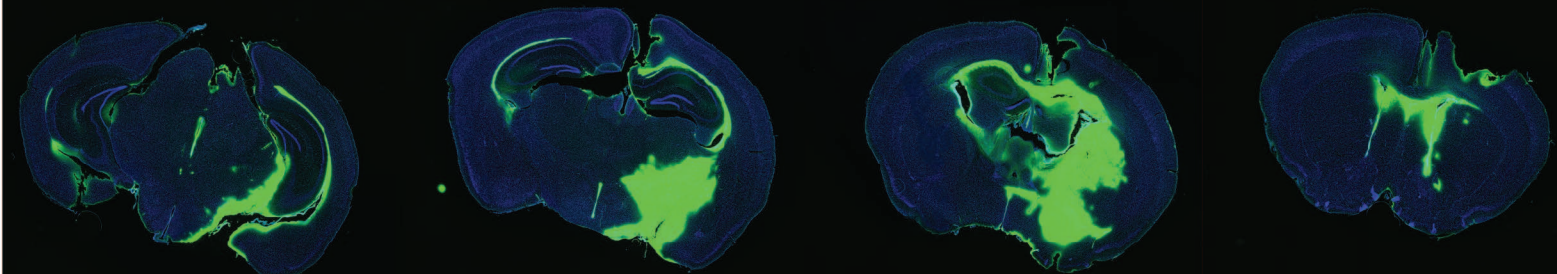

**T = 1 day**

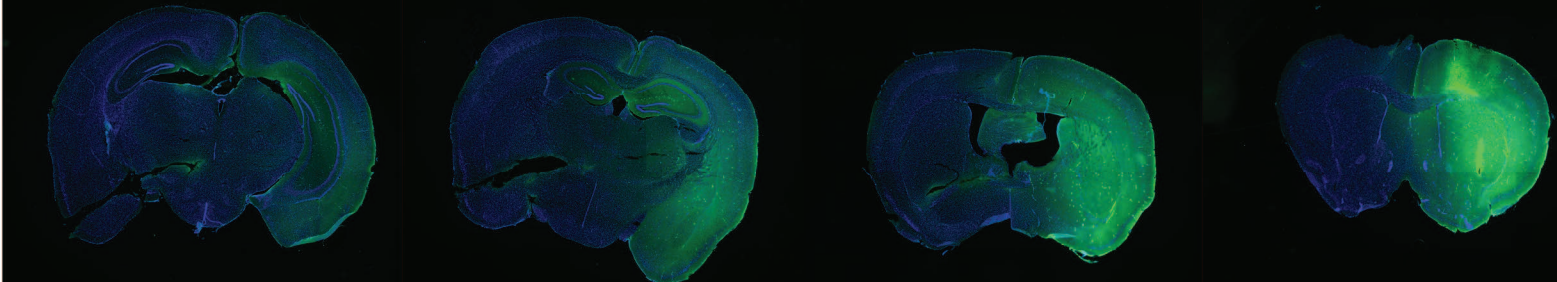

**T = 2 days**

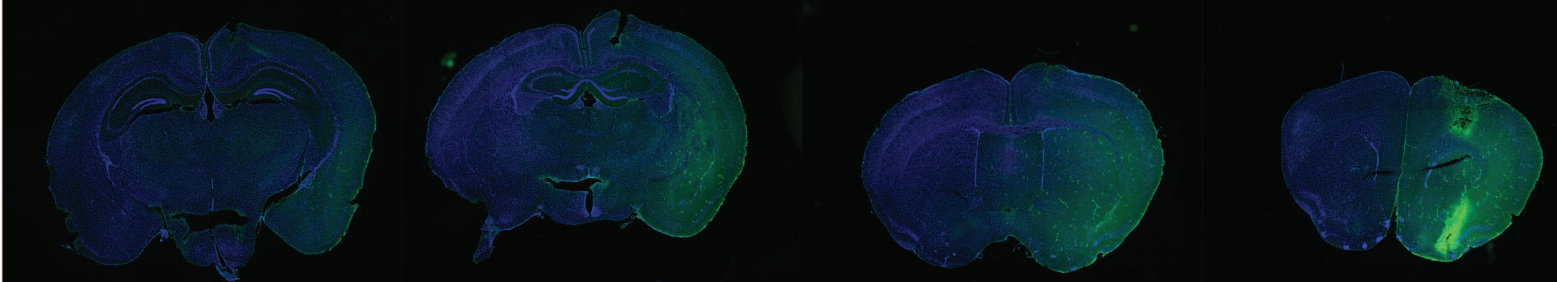

**T = 3 days**

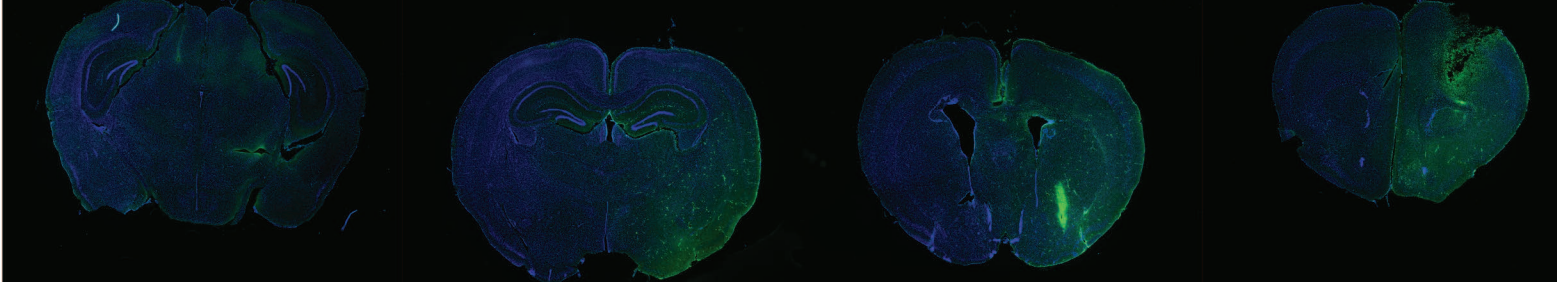

**B**

**IP Injection**

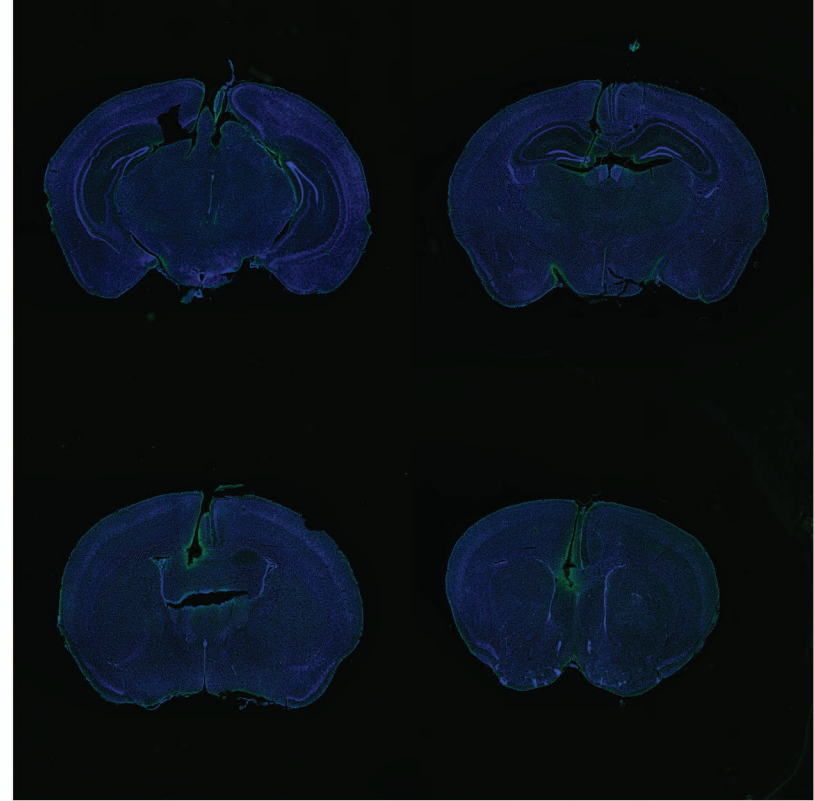

**C**

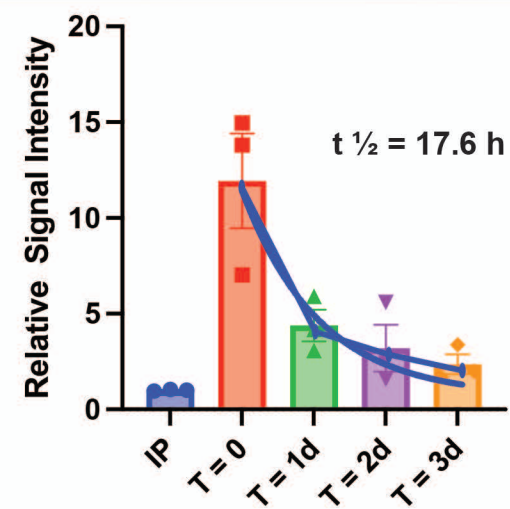

**Supplemental Figure 4. Fluorophore-labeled 1H5 allows for in vivo tracking following delivery.** (A) Following labeling of 488-1H5, it was delivered via CED in the right striatum of naïve mice, demonstrating progressive clearance over time. (B) When 488-1H5 was delivered systemically (IP), minimal signal was observed in the brain. (C) Quantification of signal intensity demonstrates an 8-fold increase in signal in the brain of mice injected with 488-1H5 via CED. A one-phase decay curve fitted to the data showed a clearance half-life of 17.6 hours.

A

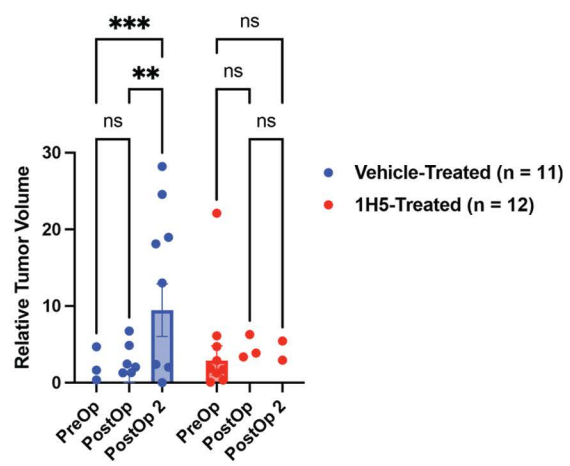

B

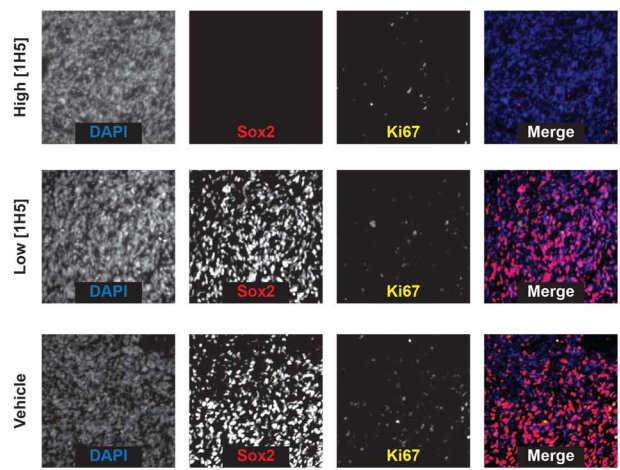

C

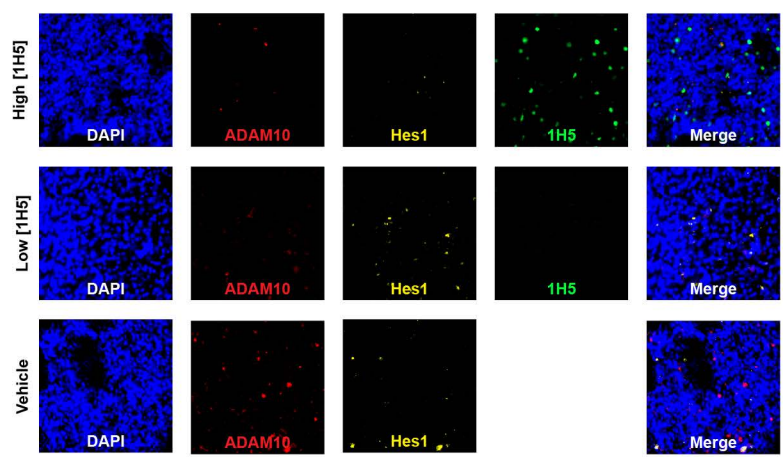

D

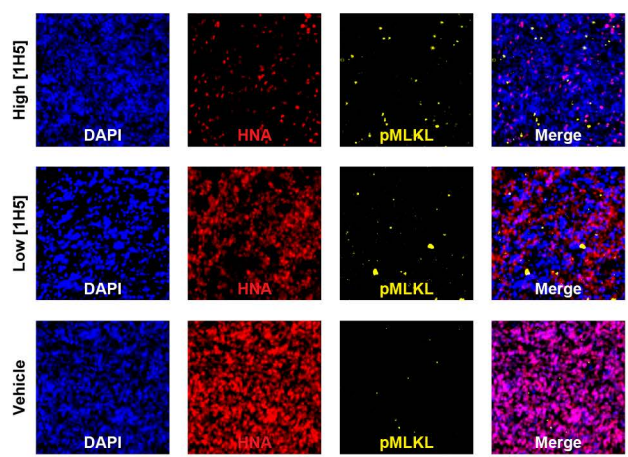

**Supplemental Figure 5. CED of 1H5 Slows Down Tumor Growth via NOTCH Inhibition in PDX Models of HGGs.** (A) Quantification of BLI signal demonstrating a significantly slower tumor growth in animals implanted with MSK-001 treated with multiple CEDs of 1H5 (n = 12) compared to vehicle (n = 11). (B-D) Immunofluorescence imaging showing a decrease in Ki67<sup>+</sup>, SOX2<sup>+</sup> (B), ADAM10<sup>+</sup>, HES1<sup>+</sup> (C), and increase in pMLKL<sup>+</sup> cells (D) following 1H5 treatment over vehicle.

| **Cell Line** | **IMPACT® Mutations** |
| --- | --- |
| **MSK-003** | EGFR, MYCN, CDKN2B, CDKN2A, ACVR1, NF1, PTEN, TERT, TRAF2, ARID1B |
| **MSK-019** | N/A |
| **MSK-018** | N/A |
| **MSK-005** | NRAS, TP53, EGFR, SOX2, DCUN1D1, MTAP, CDKN2A, CDKN2B, FOXA1, GPS2, TP53, ALOX12B, AURKB, MAP2K4, NCOR1, FLCN, DICER1, JAK2, TERT, PDE1C – EGFR, EGFR |
| **MSK-105** | MDM2, IRS2, TERT |
| **MSK-069** | AXIN1, PTEN, RB1, STAG2, TERT, TP53 |
| **MSK-201** | EGFR, MDM2, PDGFRA, KIT, KDR, CDKN2A, CDKN2B, SMYD3, ARID2, FLT1, GAB1, MLLT1, MRE11, REL, SLX4, STK19, TERT |
| **MSK-006** | N/A |
| **MSK-001** | TP53, CDKN2B, CDKN2A, BRCA1, NF1, ERG, FLT1, NUP93, PTEN, PTPRS, SF3B1, TERT, VPS8 – PRKCI |
| **MSK-023** | N/A |

**Supplemental Table 1.** List of cell lines used and their respective mutations, as determined by MSK IMPACT®.

| Antigen | Species | Ref | Vendor |
| --- | --- | --- | --- |
| β-actin | Rabbit | CST 4967 | CST |
| ADAM10 | Rabbit | 25900-1-AP | Abcam |
| ADAM10 | Mouse | MA5-38439 | R&D |
| Caspase 3 | Rabbit | CST 9662 | CST |
| Cleaved Caspase 3 | Rabbit | CST 9661 | CST |
| γH2Ax | Rabbit | CST 2595 | CST |
| Hey1 | Rabbit | PA5-40553 | Invitrogen |
| Hey2 | Rabbit | PA5-25807 | Invitrogen |
| Hes1 | Rabbit | PA5-28802 | Invitrogen |
| Hes5 | Rabbit | 22666-1-AP | ProteinTech |
| Hes6 | Rabbit | PA5-40617 | Invitrogen |
| HNA | Mouse | MAB1281 | Sigma-Aldrich |
| Ki67 | Rabbit | AB15580 | AbCam |
| MLKL | Rabbit | CST 14993 | CST |
| pMLKL | Rabbit | PA5-105678 | Invitrogen |
| Nestin | Rabbit | CST 73349 | CST |
| Nestin | Mouse | MAB5326 | Sigma-Aldrich |
| NICD Val1744 | Rabbit | CST 4147 | CST |
| NICD Val1744 | Rabbit | PA5-99448 | Invitrogen |
| NOTCH1R | Mouse | 14-5785-81 | eBioScience |
| NOTCH1R | Rabbit | MA5-32080 | Invitrogen |
| SOX2 | Rabbit | CST 3579S | CST |
| SOX2 | Rat | 149811-82 | eBioScience |
| Survivin | Rabbit | CST 2808 | CST |
| 488 | Rabbit | 710369 | Invitrogen |

**Supplemental Table 2.** Detailed information of the antibodies utilized.
